## Supplementary material for "Transcriptomic profiling of Schlemm’s canal cells reveals a lymphatic-biased identity and three major cell states": Table 1

| pathway | pval | padj | log2err | ES | NES | size | leadingEdge |
| --- | --- | --- | --- | --- | --- | --- | --- |
| REACTOME ECM PROTEOGLYCANS | 1.28E-05 | 0.0001277 | 0.59332548 | 0.6391868 | 2.54048611 | 17 | LAMA2\|TNC\|NCAM1\|TGFB2\|FN1\|COL6A1\|COMP\|MUSK\|COL2A1\|ITGB5\|COL6A2\|ITGA7\|ITGA8\|HAPLN1 |
| REACTOME INTEGRIN CELL SURFACE INTERACTIONS | 0.0040083 | 0.0200415 | 0.40701792 | 0.45224576 | 1.86792533 | 19 | ITGB8\|TNC\|COL8A1\|FN1\|CD44\|COL6A1\|COMP\|COL2A1\|ITGB5\|COL6A2\|ITGA7\|ITGA8 |
| NABA MATRISOME | 0.00731642 | 0.02438806 | 0.40701792 | 0.25408532 | 1.75462179 | 94 | LAMA2\|DPT\|ADAMTSL4\|MEGF6\|ANXA8\|CHRDL2\|PLXDC2\|MMRN1\|CHAD\|SEMA3C\|TNFAIP6\|TNC\|BMP5\|COL8A1\|IL1RN\|TGFB2\|THBS4\|FGL2\|SDC4\|GPC4\|FN1\|COL12A1\|COL6A1\|CCL11\|LGALS3\|COMP\|COL2A1 |
| REACTOME SIGNALING BY WNT | 0.02178265 | 0.05445662 | 0.35248786 | -0.4641548 | -1.696888 | 12 | SOX17\|WNT5B\|PDE6G\|SOX13\|PSMA8\|ARRB2 |
| NABA ECM GLYCOPROTEINS | 0.08726004 | 0.17452007 | 0.19991523 | 0.33346328 | 1.50424765 | 24 | LAMA2\|DPT\|MMRN1\|TNFAIP6\|TNC\|THBS4\|FGL2\|FN1\|COMP |
| WP VEGFAVEGFR2 SIGNALING PATHWAY | 0.13270142 | 0.22116904 | 0.18820415 | -0.2519388 | -1.344932 | 33 | CGNL1\|SELE\|RCAN1\|FLT1\|RND1\|MMRN2\|FMNL3\|HSPB1\|NOS3\|IGFBP7 |
| NABA MATRISOME ASSOCIATED | 0.27377049 | 0.3911007 | 0.10208011 | 0.18915585 | 1.16411213 | 59 | ADAMTSL4\|MEGF6\|ANXA8\|CHRDL2\|PLXDC2\|SEMA3C\|BMP5\|IL1RN\|TGFB2\|SDC4\|GPC4 |
| REACTOME TRANSPORT OF SMALL MOLECULES | 0.47833333 | 0.57328605 | 0.07200331 | 0.1608429 | 0.99145051 | 61 | RUNX1\|MB\|TRDN\|SLC38A3\|CASQ2\|SLC5A3\|ATP2A1\|SLC43A2\|SLC6A6\|SLC8A1\|SLC7A2\|PRKAR2B\|ABCA6\|ATP1B1\|ATP1B2\|STEAP3\|SLC7A8\|SLC18A2\|ADCY2\|SLC16A1 |
| REACTOME SLC MEDIATED TRANSMEMBRANE TRANSPORT | 0.51595745 | 0.57328605 | 0.0713053 | 0.19817622 | 0.94970303 | 28 | RUNX1\|SLC38A3\|SLC5A3\|SLC43A2\|SLC6A6\|SLC8A1\|SLC7A2\|SLC7A8\|SLC18A2\|SLC16A1 |
| REACTOME INTERLEUKIN 4 AND INTERLEUKIN 13 SIGNALING | 0.82944345 | 0.82944345 | 0.04929177 | 0.21001018 | 0.69781601 | 11 | FN1\|CCL11\|MMP3\|LIF |

|  | SEC enriched pathways |
| --- | --- |
|  | LEC enriched pathways |
