## Supplementary material for "Transcriptomic profiling of Schlemm’s canal cells reveals a lymphatic-biased identity and three major cell states": Table 2

| pathway | pval | padj | log2err | ES | NES | size | leadingEdge |
| --- | --- | --- | --- | --- | --- | --- | --- |
| REACTOME TRANSPORT OF SMALL MOLECULES | 2.50E-07 | 2.86E-06 | 0.67496286 | 0.36734478 | 2.44400992 | 112 | SLC7A5\|SLC2A1\|SLC16A1\|SLC39A10\|ATP1B2\|SLCO1C1\|SLC6A6\|SLC38A3\|SLC1A2\|GNGT1\|BSG\|ABCB1\|CA4\|TFRC\|SLC1A1\|PLTP\|CASQ2\|ATP10A\|SLC38A5\|ATP2A3\|SLC7A1\|LSR\|LRRC8C\|SLC24A1\|PRKAR2B\|AQP11\|SLC43A2\|ATP8A1\|STEAP3\|SLC30A1\|SCARB1\|SLCO2B1\|ATP1A3\|SLC16A2\|SLC11A1\|ABCA4\|SLC1A3\|RUNX1\|ANO1\|SLC31A1\|TRPC3 |
| NABA MATRISOME | 4.76E-07 | 2.86E-06 | 0.67496286 | -0.3275322 | -2.2610346 | 173 | FBLN5\|ADAMTSL1\|NTN1\|PCSK6\|ABI3BP\|MMP14\|MMRN1\|LRG1\|SEMA3D\|PRELP\|FGL2\|TIMP2\|NPNT\|SERPINE1\|THSD4\|IL7\|EGLN3\|SNED1\|IGFBP4\|SEMA3F\|ADAMTS15\|LTBP3\|INHBB\|TIMP1\|PGF\|ANXA1\|ITIH3\|COL6A3\|IGF1\|LAMA4\|COL23A1\|FBN1\|MMRN2\|S100A10\|WNT9A\|POSTN\|IL6\|PLXND1\|PDGFA\|LUM\|PAPPA\|FMOD\|AEBP1\|PLOD1\|CD109\|TGFB3\|P3H3\|ADAMTS5\|IGSF10\|EMILIN1\|ANGPTL2\|THBS1\|HGF\|FBLN2\|LAMB1\|FGF12\|COL13A1\|S100A6\|ADAMTS9\|TGFBI\|ANXA11\|FSTL1\|GDF10\|SMOC1\|LOXL2\|ITIH2\|ADAMTS4\|SERPING1 |
| REACTOME SLC MEDIATED TRANSMEMBRANE TRANSPORT | 1.09E-05 | 4.36E-05 | 0.59332548 | 0.44947394 | 2.43794772 | 49 | SLC7A5\|SLC2A1\|SLC16A1\|SLC39A10\|SLCO1C1\|SLC6A6\|SLC38A3\|SLC1A2\|BSG\|SLC1A1\|SLC38A5\|SLC7A1\|SLC24A1\|SLC43A2\|SLC30A1\|SLCO2B1\|SLC16A2\|SLC11A1\|SLC1A3\|RUNX1\|SLC31A1 |
| PID IL6 7 PATHWAY | 3.77E-05 | 0.00011317 | 0.55733224 | -0.6614673 | -2.4316911 | 16 | LBP\|FOS\|SOCS3\|JUN\|JUNB\|CEBPD\|TIMP1\|IL6\|JAK1\|MYC\|IRF1\|PTPRE\|MAPK11 |
| NABA ECM GLYCOPROTEINS | 0.00012041 | 0.00026487 | 0.5384341 | -0.4534414 | -2.3074268 | 46 | FBLN5\|NTN1\|ABI3BP\|MMRN1\|LRG1\|FGL2\|NPNT\|THSD4\|SNED1\|IGFBP4\|LTBP3\|LAMA4\|FBN1\|MMRN2\|POSTN\|AEBP1\|IGSF10\|EMILIN1\|THBS1\|FBLN2\|LAMB1 |
| REACTOME INTERLEUKIN 4 AND INTERLEUKIN 13 SIGNALING | 0.00013244 | 0.00026487 | 0.51884808 | -0.5214447 | -2.3269186 | 29 | LBP\|FOS\|SOCS3\|JUNB\|FSCN1\|CEBPD\|LCN2\|TIMP1\|ANXA1\|IL6\|CCND1\|JAK1\|HGF\|PIM1\|MYC |
| WP TGFBETA SIGNALING PATHWAY | 0.00152519 | 0.00261462 | 0.45505987 | -0.4763487 | -2.0098712 | 25 | FOS\|JUN\|FOSB\|JUNB\|ATF3\|ITGB4\|MET\|TGIF1\|DAB2\|CCND1\|THBS1\|MYC\|CAV1 |
| NABA MATRISOME ASSOCIATED | 0.00348976 | 0.00523464 | 0.4317077 | -0.2802792 | -1.7440699 | 106 | ADAMTSL1\|PCSK6\|MMP14\|SEMA3D\|TIMP2\|SERPINE1\|IL7\|EGLN3\|SEMA3F\|ADAMTS15\|INHBB\|TIMP1\|PGF\|ANXA1\|ITIH3\|IGF1\|S100A10\|WNT9A\|IL6\|PLXND1\|PDGFA\|PAPPA\|PLOD1\|CD109\|TGFB3\|P3H3\|ADAMTS5\|ANGPTL2\|HGF\|FGF12\|S100A6\|ADAMTS9\|ANXA11\|FSTL1\|GDF10\|LOXL2\|ITIH2\|ADAMTS4\|SERPING1 |
| WP VEGFAVEGFR2 SIGNALING PATHWAY | 0.0059296 | 0.00790613 | 0.40701792 | -0.3235111 | -1.7829894 | 63 | MMP14\|APOLD1\|RND1\|EGR1\|SELE\|RCAN1\|JUN\|LDB2\|NR4A1\|NFATC1\|FSCN1\|PGF\|ANXA1 |
| REACTOME INTEGRIN CELL SURFACE INTERACTIONS | 0.07641921 | 0.09170306 | 0.24133998 | 0.3350766 | 1.46690867 | 25 | BSG\|FN1\|VTN\|JAM2\|ITGB8\|VWF\|ITGB5\|ITGA1 |
| REACTOME ECM PROTEOGLYCANS | 0.57012751 | 0.62195728 | 0.06767604 | -0.2176867 | -0.9157219 | 24 | SERPINE1\|ITGA9\|COL6A3\|LAMA4\|LUM\|FMOD\|TGFB3\|LAMB1 |
| REACTOME SIGNALING BY WNT | 0.72747748 | 0.72747748 | 0.06613262 | 0.16169022 | 0.77768606 | 33 | GNGT1\|PDE6G\|LEF1\|TCF7\|PDE6B\|AXIN2\|PDE6A\|PRKCB\|GNAT2 |

|  | SEC enriched pathways |
| --- | --- |
|  | BEC enriched pathways |
