## Supplementary material for "Transcriptomic profiling of Schlemm’s canal cells reveals a lymphatic-biased identity and three major cell states": Table 3

| **Gene** | **p_val** | **avg_log2FC** | **pct.1 (B6)** | **pct.2 (129)** | **p_val_adj** |
| --- | --- | --- | --- | --- | --- |
| **Chchd10** | 3.12E-14 | -0.2872576 | 1 | 0.071 | 6.24E-11 |
| **Csf1** | 4.84E-13 | -0.3837765 | 1 | 0.089 | 9.67E-10 |
| **D830025C05Rik** | 4.84E-13 | -0.260632 | 1 | 0.089 | 9.67E-10 |
| **Pias4** | 4.84E-13 | -0.2552144 | 1 | 0.089 | 9.67E-10 |
| **Folr1** | 5.09E-13 | -0.2500469 | 0.943 | 0.071 | 1.02E-09 |
| **Atp1a2** | 6.28E-12 | -0.3964052 | 1 | 0.107 | 1.26E-08 |
| **Enpp6** | 6.28E-12 | -0.4000637 | 1 | 0.107 | 1.26E-08 |
| **Trpm1** | 1.01E-11 | -0.2824664 | 0.914 | 0.054 | 2.01E-08 |
| **Rnf168** | 1.27E-11 | -0.2789672 | 0.971 | 0.089 | 2.55E-08 |
| **Fabp4** | 2.75E-11 | -0.3127307 | 0.686 | 0.018 | 5.51E-08 |
| **Slurp1** | 2.88E-11 | -0.2741462 | 0.943 | 0.071 | 5.76E-08 |
| **Rbp1** | 6.52E-11 | -0.3004144 | 1 | 0.125 | 1.30E-07 |
| **Ephb1** | 6.52E-11 | -0.3504816 | 1 | 0.125 | 1.30E-07 |
| **Cdkn1c** | 7.69E-11 | -0.2568393 | 0.143 | 0.054 | 1.54E-07 |
| **Rasgef1b** | 1.31E-10 | -0.3253708 | 0.971 | 0.107 | 2.61E-07 |
| **Fgl2** | 2.79E-10 | -0.3496341 | 0.143 | 0.054 | 5.57E-07 |
| **Pcp4** | 6.52E-10 | -0.603055 | 1 | 0.143 | 1.30E-06 |
| **Mlana** | 6.52E-10 | -0.4861732 | 1 | 0.143 | 1.30E-06 |
| **Mnt** | 1.36E-09 | -0.2948642 | 0.171 | 0.107 | 2.72E-06 |
| **Sf3a1** | 1.43E-09 | -0.3414711 | 0.171 | 0.089 | 2.85E-06 |
| **Hmox1** | 1.70E-09 | -0.2671227 | 0.171 | 0.054 | 3.40E-06 |
| **Elmsan1** | 2.22E-08 | -0.250466 | 0.2 | 0.089 | 4.43E-05 |
| **Col23a1** | 2.56E-08 | -0.3858331 | 0.2 | 0.107 | 5.12E-05 |
| **Ifit1** | 3.45E-08 | -0.2778944 | 0.2 | 0.036 | 6.90E-05 |
| **Egr3** | 4.44E-08 | 0.37055239 | 0.2 | 0 | 8.88E-05 |
| **Gsta4** | 5.00E-08 | -0.2816064 | 0.971 | 0.161 | 0.00010009 |
| **Stc2** | 5.03E-08 | -0.3477424 | 0.714 | 0.036 | 0.00010069 |
| **Rgs9** | 1.58E-07 | -0.264223 | 0.686 | 0.036 | 0.00031527 |
| **Msi2** | 2.02E-07 | -0.2762805 | 0.229 | 0.143 | 0.00040469 |
| **Slc10a6** | 2.81E-07 | -0.4417042 | 0.229 | 0.089 | 0.00056158 |
| **Htra1** | 3.45E-07 | -0.40623 | 0.229 | 0.071 | 0.00069099 |
| **Heatr5b** | 3.45E-07 | -0.3080564 | 0.229 | 0.071 | 0.00069099 |
| **Pmel** | 4.03E-07 | -0.8024094 | 0.829 | 0.054 | 0.00080507 |
| **Msra** | 4.22E-07 | -0.2674365 | 0.229 | 0.054 | 0.00084448 |
| **Icam1** | 4.30E-07 | 0.28498753 | 0.229 | 0.089 | 0.00086067 |
| **Rdh10** | 7.39E-07 | 0.31977253 | 0.229 | 0 | 0.00147831 |
| **Naalad2** | 7.43E-07 | -0.3340133 | 0.914 | 0.143 | 0.00148504 |
| **Gpx3** | 7.88E-07 | -0.2689222 | 0.857 | 0.089 | 0.00157696 |
| **Adam12** | 2.25E-06 | -0.4548751 | 0.257 | 0.107 | 0.00450334 |
| **Slc7a1** | 3.72E-06 | -0.398215 | 0.257 | 0.071 | 0.00744012 |
| **Trmt61a** | 5.34E-06 | -0.2528762 | 0.229 | 0.018 | 0.01068584 |
| **Htr2b** | 7.14E-06 | 0.27675787 | 0.257 | 0.036 | 0.01427075 |
| **Sele** | 1.03E-05 | -0.9371836 | 1 | 0.25 | 0.02054578 |
| **Pde12** | 1.26E-05 | -0.2873676 | 0.286 | 0.179 | 0.02526023 |
| **P3h3** | 1.31E-05 | -0.2803046 | 0.286 | 0.143 | 0.0262265 |
| **Tes** | 1.85E-05 | -0.2585057 | 0.286 | 0.107 | 0.03690736 |
| **Pigo** | 2.33E-05 | -0.6520533 | 1 | 0.25 | 0.04654281 |
| **Dct** | 2.61E-05 | -0.7801816 | 1 | 0.25 | 0.05217711 |
| **Mt2** | 2.75E-05 | -0.3645836 | 0.829 | 0.107 | 0.05493549 |
| **Gpnmb** | 3.14E-05 | -0.5255118 | 0.743 | 0.036 | 0.06277992 |
| **Mta3** | 3.24E-05 | -0.3119438 | 0.286 | 0.071 | 0.0647432 |
| **Lypd2** | 7.20E-05 | -0.400059 | 0.314 | 0.143 | 0.14396681 |
| **2310001H17Rik** | 7.55E-05 | -0.6403427 | 1 | 0.268 | 0.15103289 |
| **Snhg3** | 9.16E-05 | -0.4442244 | 0.886 | 0.179 | 0.18317962 |
| **Etv3** | 9.32E-05 | 0.27300148 | 0.286 | 0 | 0.18648671 |
| **Abcb1a** | 0.00012789 | -0.337339 | 0.771 | 0.071 | 0.25577385 |
| **Cxcl2** | 0.00015779 | -0.5180705 | 0.314 | 0.125 | 0.31558375 |
| **Kmt2d** | 0.00016928 | -0.3792575 | 0.314 | 0.089 | 0.33855488 |
| **Apoe** | 0.00018447 | -0.5968615 | 1 | 0.286 | 0.36893545 |
| **Abhd17b** | 0.00025384 | -0.3429323 | 0.343 | 0.179 | 0.50768941 |
| **Itm2b** | 0.00034727 | -0.5715719 | 1 | 0.946 | 0.69454588 |
| **Dcn** | 0.00039305 | -0.2828282 | 0.857 | 0.179 | 0.7860934 |
| **Pltp** | 0.00039522 | -0.2839364 | 0.971 | 0.286 | 0.79044745 |
| **Peg3** | 0.00040664 | -0.3605774 | 0.314 | 0.036 | 0.81327022 |
| **Aqp1** | 0.0004358 | 0.49151949 | 0.314 | 0.036 | 0.87160594 |
| **Lmo4** | 0.00057168 | -0.8040746 | 0.4 | 0.268 | 1 |
| **Zfp46** | 0.00068986 | -0.2712024 | 0.314 | 0.054 | 1 |
| **Nkd1** | 0.00069985 | -0.3555677 | 0.343 | 0.107 | 1 |
| **Plaur** | 0.00070943 | 0.44108998 | 0.314 | 0 | 1 |
| **Lst1** | 0.00093735 | -0.3601317 | 0.371 | 0.179 | 1 |
| **Tgfbi** | 0.00095788 | -0.3684122 | 0.343 | 0.089 | 1 |
| **Sec22b** | 0.00102905 | -0.3606681 | 0.371 | 0.179 | 1 |
| **Myc** | 0.00106142 | -0.3872047 | 0.371 | 0.179 | 1 |
| **Nbl1** | 0.00124431 | -0.4457405 | 0.371 | 0.161 | 1 |
| **Oat** | 0.00130243 | -0.4844033 | 0.343 | 0.071 | 1 |
| **Lncpint** | 0.00134408 | -0.3049655 | 0.343 | 0.071 | 1 |
| **5031425E22Rik** | 0.00136397 | -0.2824938 | 0.371 | 0.161 | 1 |
| **Ccl21a** | 0.00143282 | -2.2368647 | 0.4 | 0.268 | 1 |
| **H2-Aa** | 0.00165173 | 0.28460315 | 0.686 | 0.018 | 1 |
| **Ntn4** | 0.00169237 | -0.3615339 | 0.371 | 0.143 | 1 |
| **Pcdh17** | 0.00179178 | -0.3167384 | 0.4 | 0.25 | 1 |
| **Krt5** | 0.00190811 | -0.3449657 | 0.371 | 0.143 | 1 |
| **Timp3** | 0.0022855 | -0.8761295 | 1 | 0.786 | 1 |
| **S100a1** | 0.00251646 | -0.4709522 | 0.971 | 0.304 | 1 |
| **Cldn5** | 0.00300967 | -0.8027568 | 0.971 | 0.839 | 1 |
| **Akap12** | 0.00334552 | 0.54852886 | 0.343 | 0.018 | 1 |
| **Flt4** | 0.00471508 | -0.2929918 | 0.943 | 0.304 | 1 |
| **Rpl36a** | 0.00635003 | -0.5661309 | 1 | 0.893 | 1 |
| **Sik1** | 0.00697705 | -0.4085242 | 0.4 | 0.143 | 1 |
| **A830019P07Rik** | 0.00726262 | -0.2603155 | 0.343 | 0.054 | 1 |
| **Mt1** | 0.00727901 | -0.4826229 | 1 | 0.375 | 1 |
| **Timp1** | 0.00772847 | 0.51953261 | 0.371 | 0.071 | 1 |
| **Tiparp** | 0.00775366 | -0.302615 | 0.4 | 0.143 | 1 |
| **Zfp574** | 0.00815868 | -0.2569245 | 0.371 | 0.054 | 1 |
| **Dennd4a** | 0.00825022 | -0.6752471 | 0.457 | 0.25 | 1 |
| **Slc45a3** | 0.00838095 | -0.270281 | 0.371 | 0.054 | 1 |
| **Klf4** | 0.01029332 | -0.5924954 | 0.971 | 0.911 | 1 |
| **mt-Nd5** | 0.01152139 | -0.645719 | 0.943 | 0.821 | 1 |
| **Slfn3** | 0.01193373 | -0.3045298 | 1 | 0.357 | 1 |
| **Plekhg3** | 0.01297636 | -0.330813 | 0.343 | 0.018 | 1 |
| **Rpl22l1** | 0.01328726 | -0.4549634 | 1 | 0.929 | 1 |
| **Il17ra** | 0.01351736 | -0.4184936 | 0.457 | 0.25 | 1 |
| **Stt3b** | 0.01360177 | -1.0792393 | 0.571 | 0.393 | 1 |
| **Serpine1** | 0.01397551 | -0.2871424 | 0.429 | 0.196 | 1 |
| **Wnt2** | 0.01451392 | 0.25867188 | 0.371 | 0.018 | 1 |
| **Exoc3l2** | 0.01488995 | 0.25485812 | 0.371 | 0.018 | 1 |
| **Sfxn2** | 0.01546244 | -0.3851957 | 0.371 | 0.054 | 1 |
| **Fah** | 0.01568922 | -0.2868699 | 0.714 | 0.107 | 1 |
| **Firre** | 0.01646539 | -0.3127052 | 0.657 | 0.036 | 1 |
| **Il6** | 0.01703408 | -0.4298634 | 0.4 | 0.089 | 1 |
| **Zfp593** | 0.01703408 | -0.3988337 | 0.4 | 0.089 | 1 |
| **Eln** | 0.01708519 | -0.3997895 | 0.829 | 0.232 | 1 |
| **Rpl37a** | 0.01819335 | -0.3731844 | 1 | 1 | 1 |
| **Gch1** | 0.01914699 | 0.40344106 | 0.371 | 0 | 1 |
| **Iigp1** | 0.01997871 | 0.41147247 | 0.4 | 0.107 | 1 |
| **Ivns1abp** | 0.02192288 | -0.2989903 | 0.429 | 0.143 | 1 |
| **Sema3f** | 0.02264758 | -0.7455812 | 0.686 | 0.554 | 1 |
| **Spry1** | 0.02404706 | -0.3626365 | 0.429 | 0.143 | 1 |
| **Trim35** | 0.02404706 | -0.2634841 | 0.429 | 0.143 | 1 |
| **Ipo11** | 0.02568386 | -0.3753476 | 0.857 | 0.304 | 1 |
| **Duox2** | 0.02802923 | -0.5542768 | 0.457 | 0.196 | 1 |
| **Tbx1** | 0.02865674 | -0.4939604 | 0.457 | 0.196 | 1 |
| **Mylk** | 0.03328954 | -0.5129122 | 0.457 | 0.179 | 1 |
| **Gm1673** | 0.03552271 | -0.2746482 | 0.457 | 0.179 | 1 |
| **Nr4a2** | 0.03798004 | -0.4691263 | 0.457 | 0.196 | 1 |
| **Klk8** | 0.038396 | -0.2512415 | 0.914 | 0.375 | 1 |
| **Rcan1** | 0.03993542 | -0.8972014 | 0.771 | 0.643 | 1 |
| **Zfand2a** | 0.04037384 | 0.26829478 | 0.4 | 0.036 | 1 |
| **Slc41a1** | 0.04037808 | -0.3631653 | 0.429 | 0.107 | 1 |
| **Entpd1** | 0.04039311 | -0.3805628 | 0.829 | 0.268 | 1 |
| **mt-Nd4l** | 0.0409571 | -0.6667788 | 1 | 0.875 | 1 |
| **Fam189a2** | 0.04477441 | -0.6440195 | 0.6 | 0.393 | 1 |
| **Apold1** | 0.04951049 | -0.6984794 | 0.943 | 0.786 | 1 |
| **Nsg1** | 0.05030957 | -0.6020614 | 0.971 | 0.75 | 1 |
| **Ier5l** | 0.0505164 | -0.5819718 | 0.514 | 0.268 | 1 |
| **Pim3** | 0.05054423 | -0.849765 | 0.571 | 0.339 | 1 |
| **Cobll1** | 0.05127578 | -0.3099875 | 0.657 | 0.071 | 1 |
| **Fxyd6** | 0.05304615 | -0.7132838 | 0.971 | 0.714 | 1 |
| **1810026B05Rik** | 0.05612879 | -0.4234821 | 0.743 | 0.179 | 1 |
| **Anxa2** | 0.05879093 | -0.407991 | 1 | 0.821 | 1 |
| **Rapgef5** | 0.05976832 | -0.4224213 | 1 | 0.786 | 1 |
| **Gngt2** | 0.06105577 | -0.4552323 | 1 | 0.839 | 1 |
| **Lmna** | 0.06802381 | -0.4349872 | 1 | 0.804 | 1 |
| **Rdx** | 0.07289427 | -0.84679 | 0.886 | 0.625 | 1 |
| **Gadd45a** | 0.07404143 | -0.3989594 | 0.429 | 0.071 | 1 |
| **Lgals1** | 0.07795174 | -0.2593598 | 0.686 | 0.125 | 1 |
| **Mfsd4a** | 0.08192468 | -0.3275749 | 0.514 | 0.25 | 1 |
| **Timp2** | 0.08299191 | -0.3373508 | 1 | 0.929 | 1 |
| **Cd55** | 0.0857329 | -0.3368075 | 0.743 | 0.196 | 1 |
| **Adamtsl1** | 0.08881211 | -0.3692159 | 0.971 | 0.839 | 1 |
| **Dnaja1** | 0.09059502 | -0.7742671 | 0.943 | 0.661 | 1 |
| **Arl4c** | 0.09074231 | -0.3668896 | 0.486 | 0.179 | 1 |
| **Plec** | 0.09259129 | -0.5855566 | 0.971 | 0.696 | 1 |
| **Ablim1** | 0.09292914 | 0.25083523 | 0.429 | 0.054 | 1 |
| **Prox1** | 0.09362077 | -0.3648119 | 0.886 | 0.339 | 1 |
| **Selenop** | 0.09436896 | -0.874302 | 0.943 | 0.714 | 1 |
| **Ift74** | 0.0983664 | 0.3215848 | 0.429 | 0.054 | 1 |
| **Trp53i11** | 0.10115464 | -0.5878966 | 0.971 | 0.732 | 1 |
| **Ifi27l2a** | 0.10129793 | -1.0931468 | 0.914 | 0.75 | 1 |
| **Ptgds** | 0.10460446 | -0.8863597 | 1 | 0.732 | 1 |
| **F2r** | 0.10793359 | -0.5724008 | 1 | 0.464 | 1 |
| **F8** | 0.11459905 | -0.3129879 | 0.714 | 0.179 | 1 |
| **Ppl** | 0.11891227 | -0.2963306 | 0.486 | 0.161 | 1 |
| **Large1** | 0.12003544 | -0.2638181 | 0.429 | 0.036 | 1 |
| **Dync1i2** | 0.12965733 | -0.4566815 | 1 | 0.768 | 1 |
| **Btbd3** | 0.13408746 | -0.7702418 | 0.743 | 0.536 | 1 |
| **Aph1b** | 0.13678325 | -0.2580309 | 0.629 | 0.071 | 1 |
| **Sac3d1** | 0.13918509 | -0.5683627 | 0.629 | 0.071 | 1 |
| **Cebpd** | 0.14086289 | -0.3326214 | 0.971 | 0.911 | 1 |
| **Cd300lg** | 0.1426388 | -0.6492774 | 0.514 | 0.196 | 1 |
| **Plce1** | 0.14876588 | -0.4528443 | 0.486 | 0.143 | 1 |
| **Cp** | 0.14900405 | -0.5049171 | 0.943 | 0.732 | 1 |
| **Pard6g** | 0.15258275 | -0.4936582 | 0.657 | 0.429 | 1 |
| **2200002D01Rik** | 0.1558463 | -0.3314662 | 0.886 | 0.357 | 1 |
| **Rps27** | 0.15696179 | -0.3131628 | 1 | 1 | 1 |
| **Ier2** | 0.15793845 | -0.8369669 | 0.829 | 0.571 | 1 |
| **Klf2** | 0.15936456 | -0.4408619 | 1 | 0.964 | 1 |
| **Rhob** | 0.16787527 | -0.5049318 | 0.829 | 0.696 | 1 |
| **Nts** | 0.16871925 | -0.9221373 | 1 | 0.518 | 1 |
| **Sox7** | 0.16989811 | -0.4558229 | 0.6 | 0.339 | 1 |
| **Pkhd1l1** | 0.17110989 | -0.3581793 | 0.6 | 0.357 | 1 |
| **Btg1** | 0.17692111 | -0.3936313 | 0.971 | 0.893 | 1 |
| **Gem** | 0.17923628 | 0.2989028 | 0.743 | 0.286 | 1 |
| **Olfml2a** | 0.1879126 | -0.5283813 | 0.714 | 0.464 | 1 |
| **Hspb1** | 0.18978501 | -0.9446772 | 0.943 | 0.768 | 1 |
| **Rgs16** | 0.1900378 | -0.572107 | 0.943 | 0.804 | 1 |
| **S100a11** | 0.19013674 | -0.3451405 | 1 | 0.821 | 1 |
| **Dusp1** | 0.19148278 | -0.6477607 | 0.914 | 0.679 | 1 |
| **Nfatc1** | 0.19633918 | -0.282312 | 0.486 | 0.125 | 1 |
| **Cryab** | 0.19637437 | -0.4697135 | 0.829 | 0.321 | 1 |
| **Myzap** | 0.20019322 | -0.4414121 | 0.629 | 0.375 | 1 |
| **Upk3bl** | 0.20350838 | -0.4077241 | 0.629 | 0.089 | 1 |
| **Tgm2** | 0.20797453 | -0.8025689 | 0.629 | 0.357 | 1 |
| **Arl4a** | 0.20872867 | -0.7113721 | 0.943 | 0.679 | 1 |
| **Ltbp4** | 0.21870956 | -0.4538504 | 1 | 0.75 | 1 |
| **Pdgfa** | 0.2240168 | -0.4879394 | 0.8 | 0.554 | 1 |
| **Fosb** | 0.22520518 | -0.2934584 | 0.829 | 0.786 | 1 |
| **Tgfb1** | 0.23459665 | -0.5681138 | 0.657 | 0.375 | 1 |
| **Dst** | 0.23792794 | -0.6741275 | 0.886 | 0.589 | 1 |
| **Hbegf** | 0.24192766 | 0.35015689 | 0.457 | 0.054 | 1 |
| **Lamb1** | 0.24754045 | -0.4654346 | 0.543 | 0.214 | 1 |
| **Ppp3ca** | 0.25451227 | -0.5671294 | 0.829 | 0.321 | 1 |
| **Cd74** | 0.25689679 | 0.48914986 | 0.457 | 0.054 | 1 |
| **Akap13** | 0.25759184 | -0.7411 | 0.914 | 0.571 | 1 |
| **Sparc** | 0.26544347 | -0.3457959 | 1 | 0.911 | 1 |
| **Trf** | 0.26592252 | -0.3834783 | 0.743 | 0.25 | 1 |
| **Rhoc** | 0.26868426 | -0.3426185 | 0.971 | 0.821 | 1 |
| **Cd302** | 0.26933248 | -0.4484141 | 0.657 | 0.375 | 1 |
| **Cdkn1a** | 0.26979078 | -0.7014192 | 0.8 | 0.518 | 1 |
| **Fcer1g** | 0.27155579 | -0.5970249 | 0.657 | 0.143 | 1 |
| **Sat1** | 0.27316851 | -1.4273912 | 1 | 0.625 | 1 |
| **Ugcg** | 0.27406679 | -0.2763491 | 0.514 | 0.161 | 1 |
| **Aldh1a3** | 0.27645113 | -0.5404077 | 0.686 | 0.179 | 1 |
| **Ccn1** | 0.28416896 | -0.5911824 | 0.829 | 0.375 | 1 |
| **Meox1** | 0.28509368 | -0.7650324 | 0.857 | 0.536 | 1 |
| **Plcg2** | 0.28581095 | -0.2874994 | 0.629 | 0.107 | 1 |
| **Cfap298** | 0.28581095 | -0.2685153 | 0.629 | 0.107 | 1 |
| **Ufsp2** | 0.29847459 | -0.5355177 | 0.543 | 0.196 | 1 |
| **Isg15** | 0.30469081 | -0.9405508 | 0.686 | 0.393 | 1 |
| **Rnf19b** | 0.31617778 | -0.405126 | 0.514 | 0.143 | 1 |
| **Myadm** | 0.31765363 | -0.7273949 | 0.8 | 0.482 | 1 |
| **Gadd45g** | 0.32075377 | -0.375077 | 0.829 | 0.393 | 1 |
| **Nfkbia** | 0.32137483 | -0.3485698 | 0.971 | 0.839 | 1 |
| **AA467197** | 0.32335545 | -0.8094808 | 0.571 | 0.232 | 1 |
| **Thbd** | 0.32570616 | -0.5384821 | 0.857 | 0.589 | 1 |
| **Ccl2** | 0.33249232 | 0.79309784 | 0.486 | 0.107 | 1 |
| **Flrt2** | 0.33473314 | -0.3908973 | 0.6 | 0.304 | 1 |
| **Bok** | 0.33479036 | -0.4854477 | 0.771 | 0.5 | 1 |
| **Synpo** | 0.34055052 | -0.344931 | 0.629 | 0.321 | 1 |
| **Nme2** | 0.36304113 | 0.36752088 | 1 | 0.982 | 1 |
| **Nudt4** | 0.3655962 | -0.4328306 | 0.543 | 0.179 | 1 |
| **Galnt1** | 0.38759943 | -0.334896 | 0.943 | 0.679 | 1 |
| **Sparcl1** | 0.38768307 | -0.4043222 | 1 | 0.518 | 1 |
| **Banp** | 0.38770717 | -0.3477732 | 0.486 | 0.071 | 1 |
| **Pthlh** | 0.38877312 | -0.6242748 | 0.571 | 0.214 | 1 |
| **Neat1** | 0.39224288 | -0.7449476 | 0.857 | 0.518 | 1 |
| **Mxra8** | 0.39597552 | -0.3443693 | 0.914 | 0.643 | 1 |
| **Ctsl** | 0.40444481 | -0.4392868 | 0.886 | 0.393 | 1 |
| **Ntn1** | 0.40678269 | -0.6801825 | 0.829 | 0.536 | 1 |
| **Fos** | 0.4076583 | -0.2542851 | 1 | 0.946 | 1 |
| **Sema3a** | 0.40800992 | -0.2541704 | 0.829 | 0.375 | 1 |
| **Map2k3** | 0.42555752 | 0.32760821 | 0.571 | 0.054 | 1 |
| **Dleu2** | 0.42914842 | -0.4693607 | 1 | 0.5 | 1 |
| **Fscn1** | 0.43445218 | -0.6133105 | 0.743 | 0.411 | 1 |
| **Socs3** | 0.43527694 | -0.3605178 | 0.886 | 0.768 | 1 |
| **Prox1os** | 0.43669832 | 0.38479349 | 0.543 | 0 | 1 |
| **Gpha2** | 0.44702105 | 0.28166286 | 0.571 | 0.054 | 1 |
| **Ifrd1** | 0.44827235 | -0.4657615 | 0.686 | 0.393 | 1 |
| **Calcrl** | 0.45123059 | -0.4527294 | 0.857 | 0.571 | 1 |
| **Slco2a1** | 0.45603854 | -0.6544387 | 0.686 | 0.357 | 1 |
| **Procr** | 0.46952237 | -0.4852449 | 0.571 | 0.214 | 1 |
| **Gpm6a** | 0.4701945 | -0.3324038 | 0.657 | 0.339 | 1 |
| **Ptx3** | 0.47469656 | -0.2628204 | 0.486 | 0.054 | 1 |
| **Sqstm1** | 0.47665303 | -0.5345336 | 0.829 | 0.357 | 1 |
| **Bcr** | 0.48092983 | -0.6832728 | 0.829 | 0.429 | 1 |
| **Mgll** | 0.48708332 | 0.30764085 | 0.629 | 0.179 | 1 |
| **Mmrn1** | 0.48921677 | -0.4873483 | 0.714 | 0.25 | 1 |
| **Bambi** | 0.49241137 | -0.7179657 | 0.629 | 0.286 | 1 |
| **Bin1** | 0.49467791 | -0.5704058 | 0.857 | 0.393 | 1 |
| **Tmx4** | 0.50007383 | -0.2830074 | 0.6 | 0.25 | 1 |
| **Clu** | 0.50577346 | -0.3837207 | 0.943 | 0.518 | 1 |
| **Egr1** | 0.51575308 | -0.2513094 | 0.886 | 0.5 | 1 |
| **Lbp** | 0.51896304 | -0.289937 | 0.914 | 0.607 | 1 |
| **Actn1** | 0.52522464 | -0.3984874 | 0.629 | 0.286 | 1 |
| **Gm13889** | 0.54732097 | -0.5293781 | 0.629 | 0.161 | 1 |
| **Gdf15** | 0.55113877 | -0.3167669 | 0.543 | 0.018 | 1 |
| **Dnajb1** | 0.5511856 | -0.3557925 | 0.571 | 0.196 | 1 |
| **Anxa1** | 0.56394163 | -0.5997482 | 0.971 | 0.696 | 1 |
| **Fbln5** | 0.56502466 | -0.3400475 | 0.943 | 0.5 | 1 |
| **Vwa1** | 0.57059691 | 0.28600773 | 0.514 | 0.107 | 1 |
| **Prcp** | 0.57178309 | -0.4878096 | 0.886 | 0.536 | 1 |
| **Rassf9** | 0.57342659 | -0.7622664 | 0.8 | 0.429 | 1 |
| **Maf** | 0.57686355 | -0.4391527 | 0.971 | 0.518 | 1 |
| **Abi3bp** | 0.58633273 | -0.5278899 | 0.743 | 0.464 | 1 |
| **Clic5** | 0.58873908 | -0.3277009 | 0.829 | 0.536 | 1 |
| **Ppp1r15a** | 0.59757107 | -0.4283742 | 0.771 | 0.339 | 1 |
| **Ptpn1** | 0.60235562 | -0.4869062 | 0.771 | 0.321 | 1 |
| **Tob1** | 0.61076663 | -0.2855743 | 0.657 | 0.196 | 1 |
| **Cav1** | 0.61969348 | -0.5972243 | 0.886 | 0.429 | 1 |
| **Nupr1** | 0.62737642 | -0.299297 | 1 | 0.911 | 1 |
| **Hs3st1** | 0.64210421 | -0.3779462 | 0.629 | 0.286 | 1 |
| **Itga5** | 0.65383271 | -0.4124025 | 0.6 | 0.125 | 1 |
| **Tnfaip3** | 0.65643997 | 0.47254193 | 0.543 | 0.036 | 1 |
| **Itm2c** | 0.66211345 | -0.7102068 | 0.914 | 0.446 | 1 |
| **Smad1** | 0.66622541 | -0.3658688 | 0.914 | 0.607 | 1 |
| **Filip1l** | 0.66669819 | -0.3464784 | 0.543 | 0.125 | 1 |
| **Tssc4** | 0.66669819 | -0.3494788 | 0.543 | 0.125 | 1 |
| **Myo1b** | 0.66810023 | -0.4410188 | 0.6 | 0.214 | 1 |
| **Gm19705** | 0.67059384 | -0.5382606 | 0.714 | 0.357 | 1 |
| **Ppfibp1** | 0.67719285 | -0.4733294 | 0.943 | 0.554 | 1 |
| **Rgs2** | 0.6779773 | -0.5846742 | 0.629 | 0.179 | 1 |
| **Nfkbiz** | 0.68371393 | -0.4373373 | 0.714 | 0.375 | 1 |
| **Nudt3** | 0.68618058 | -0.3740969 | 0.543 | 0.125 | 1 |
| **Prelp** | 0.68989291 | -0.5759431 | 0.714 | 0.375 | 1 |
| **Pdlim1** | 0.69681538 | -0.7611792 | 0.771 | 0.393 | 1 |
| **Hspa1b** | 0.70148046 | 0.60097259 | 0.8 | 0.554 | 1 |
| **Hspa1a** | 0.70430776 | 0.61496092 | 0.714 | 0.429 | 1 |
| **Comt** | 0.70528156 | -0.7022772 | 0.743 | 0.304 | 1 |
| **Rps21** | 0.71659111 | -0.2579884 | 1 | 0.982 | 1 |
| **Cda** | 0.71673182 | -0.2717374 | 0.743 | 0.429 | 1 |
| **Txnip** | 0.71874328 | -0.5215417 | 0.857 | 0.5 | 1 |
| **Tlr2** | 0.7226083 | -0.3756613 | 0.571 | 0.089 | 1 |
| **Myh9** | 0.72738937 | -0.6345111 | 0.8 | 0.375 | 1 |
| **Slco3a1** | 0.72789634 | -0.2713441 | 0.629 | 0.25 | 1 |
| **Odc1** | 0.72976677 | -0.5313562 | 0.771 | 0.446 | 1 |
| **3300005D01Rik** | 0.74129651 | 0.31503722 | 0.514 | 0.071 | 1 |
| **Mast4** | 0.74504156 | -0.5154869 | 0.771 | 0.339 | 1 |
| **Man1a** | 0.74532052 | -0.3003105 | 0.6 | 0.214 | 1 |
| **Emp1** | 0.75822876 | -0.3945813 | 0.971 | 0.625 | 1 |
| **Irf1** | 0.75973175 | -0.8512278 | 0.8 | 0.393 | 1 |
| **Sned1** | 0.76287268 | -0.3069187 | 0.714 | 0.304 | 1 |
| **Gm11837** | 0.76296892 | -0.3603038 | 0.629 | 0.179 | 1 |
| **Dusp8** | 0.77410112 | -0.2855348 | 0.6 | 0.143 | 1 |
| **Lrg1** | 0.7881966 | -0.8404555 | 0.857 | 0.518 | 1 |
| **Crip1** | 0.7896489 | -0.2636357 | 0.943 | 0.625 | 1 |
| **Hist1h1c** | 0.79217321 | -0.2579032 | 0.543 | 0.107 | 1 |
| **Tkt** | 0.79906466 | -0.4099295 | 0.543 | 0.107 | 1 |
| **Marcksl1** | 0.79918802 | -0.4563048 | 0.829 | 0.429 | 1 |
| **Dnm1l** | 0.80203271 | -0.2565585 | 0.714 | 0.304 | 1 |
| **Igfbp5** | 0.80973514 | 0.49326111 | 0.943 | 0.839 | 1 |
| **Klf6** | 0.82156332 | -0.4302772 | 0.971 | 0.643 | 1 |
| **Slfn2** | 0.82180767 | -0.2880962 | 0.714 | 0.304 | 1 |
| **Plat** | 0.82785304 | -0.4168399 | 0.686 | 0.286 | 1 |
| **Sgk1** | 0.83039093 | -0.6566155 | 0.771 | 0.375 | 1 |
| **Sntb2** | 0.83279684 | -0.5235489 | 0.886 | 0.5 | 1 |
| **Hspe1** | 0.83606024 | -0.3073444 | 0.743 | 0.339 | 1 |
| **Itga9** | 0.83840665 | -0.2643468 | 0.829 | 0.446 | 1 |
| **Tpm1** | 0.84060522 | -0.3277267 | 0.686 | 0.268 | 1 |
| **Zfp36l2** | 0.84266238 | -0.2667037 | 0.714 | 0.339 | 1 |
| **Grina** | 0.84310195 | -0.6125456 | 0.743 | 0.357 | 1 |
| **Adamts5** | 0.85081038 | -0.5377075 | 0.8 | 0.411 | 1 |
| **Emp2** | 0.85288244 | -0.3583304 | 0.657 | 0.232 | 1 |
| **Tjp1** | 0.85630288 | -0.2899932 | 0.771 | 0.357 | 1 |
| **Adamts1** | 0.85703619 | -0.5758147 | 0.743 | 0.393 | 1 |
| **Csrp1** | 0.85767874 | -0.5980636 | 0.857 | 0.429 | 1 |
| **Taf1d** | 0.85965237 | -0.2546848 | 0.657 | 0.232 | 1 |
| **Serping1** | 0.86173098 | -0.6520791 | 0.571 | 0.107 | 1 |
| **Nfatc2** | 0.86173098 | -0.3711466 | 0.571 | 0.107 | 1 |
| **Ccdc88c** | 0.8632734 | -0.4020845 | 0.571 | 0.143 | 1 |
| **Nectin2** | 0.89446764 | -0.342757 | 0.657 | 0.268 | 1 |
| **Cdh13** | 0.89552773 | -0.3788631 | 0.714 | 0.321 | 1 |
| **Ppp1r14b** | 0.89804856 | -0.6373514 | 0.914 | 0.518 | 1 |
| **Capg** | 0.90223287 | -0.3167029 | 0.714 | 0.321 | 1 |
| **Wnk1** | 0.9105295 | -0.6297211 | 0.857 | 0.446 | 1 |
| **Plpp3** | 0.91729081 | -0.5051918 | 0.886 | 0.464 | 1 |
| **Tns3** | 0.92270087 | 0.26524388 | 0.514 | 0.018 | 1 |
| **Tsc22d1** | 0.92374044 | -0.4137756 | 0.829 | 0.446 | 1 |
| **Cd63** | 0.92426567 | -0.4824916 | 0.943 | 0.518 | 1 |
| **Cavin2** | 0.92479593 | -0.2657029 | 0.971 | 0.625 | 1 |
| **Pde4b** | 0.92753358 | -0.3976073 | 0.629 | 0.214 | 1 |
| **Bhlhe40** | 0.93110761 | -0.5097619 | 0.886 | 0.571 | 1 |
| **Ly6c1** | 0.93125458 | -0.4970145 | 0.971 | 0.607 | 1 |
| **Pkig** | 0.93655509 | -0.2510904 | 0.743 | 0.393 | 1 |
| **Zmiz1** | 0.94862737 | -0.2655868 | 0.657 | 0.25 | 1 |
| **Hjurp** | 0.95701191 | -0.465158 | 0.886 | 0.482 | 1 |
| **Hif1a** | 0.9638369 | -0.4965893 | 0.943 | 0.554 | 1 |
| **Ebf1** | 0.96830002 | 0.30076352 | 0.571 | 0.143 | 1 |
| **Sox4** | 0.96927928 | -0.2687767 | 0.657 | 0.268 | 1 |
| **Kdr** | 0.97044467 | -0.5465946 | 0.943 | 0.571 | 1 |
| **Ddx3y** | 0.97559212 | -0.3196767 | 0.6 | 0.179 | 1 |
| **Cd109** | 0.97634514 | -0.2705965 | 0.714 | 0.321 | 1 |
| **Sema3d** | 0.97671049 | -0.5204789 | 0.829 | 0.429 | 1 |
| **Plk2** | 0.98355769 | -0.549535 | 0.914 | 0.554 | 1 |
| **Ier3** | 0.99017645 | -0.9875788 | 1 | 0.625 | 1 |
| **Il6st** | 0.99654447 | -0.358729 | 0.629 | 0.214 | 1 |
| **Hes1** | 0.99673964 | -0.3514782 | 0.971 | 0.75 | 1 |
