## Supplementary material for "Transcriptomic profiling of Schlemm’s canal cells reveals a lymphatic-biased identity and three major cell states": Table 5

| **SYMBOL** | **p_val** | **avg_log2FC** | **pct.1** | **pct.2** | **p_val_adj** |
| --- | --- | --- | --- | --- | --- |
| **Npnt** | 1.39E-18 | 1.58982976 | 0.89 | 0.262 | 2.86E-14 |
| **Klk8** | 6.63E-14 | 1.41154383 | 0.44 | 0.055 | 1.37E-09 |
| **Myadm** | 1.63E-13 | 0.89967055 | 0.593 | 0.131 | 3.37E-09 |
| **Slco2a1** | 3.86E-13 | 1.04844649 | 0.495 | 0.087 | 7.95E-09 |
| **Anxa1** | 4.55E-13 | 1.03185573 | 0.802 | 0.268 | 9.36E-09 |
| **Ly6c1** | 4.83E-13 | 0.91651258 | 0.648 | 0.164 | 9.95E-09 |
| **Cd109** | 1.13E-12 | 0.79876857 | 0.396 | 0.049 | 2.33E-08 |
| **Mxra7** | 1.14E-12 | 0.66714606 | 0.418 | 0.055 | 2.36E-08 |
| **Adamts5** | 6.03E-12 | 0.76703168 | 0.56 | 0.126 | 1.24E-07 |
| **1500009L16Rik** | 6.17E-12 | 0.90450542 | 0.516 | 0.115 | 1.27E-07 |
| **Smad4** | 6.32E-12 | 0.68678228 | 0.451 | 0.082 | 1.30E-07 |
| **Ly6a** | 9.38E-12 | 0.98853893 | 0.582 | 0.153 | 1.93E-07 |
| **Ccnd2** | 1.21E-11 | 0.57058629 | 0.692 | 0.191 | 2.50E-07 |
| **Mknk2** | 4.19E-11 | 0.91860272 | 0.451 | 0.098 | 8.64E-07 |
| **Ackr3** | 7.55E-11 | 0.71665776 | 0.495 | 0.109 | 1.55E-06 |
| **Phactr4** | 1.28E-10 | 0.69339351 | 0.341 | 0.044 | 2.64E-06 |
| **Nts** | 2.23E-10 | 1.53798611 | 0.538 | 0.153 | 4.59E-06 |
| **Hexa** | 3.42E-10 | 0.68291229 | 0.473 | 0.109 | 7.04E-06 |
| **Hmgn2** | 4.25E-10 | 0.5521214 | 0.462 | 0.109 | 8.75E-06 |
| **Dpysl3** | 5.94E-10 | 0.48808045 | 0.451 | 0.093 | 1.22E-05 |
| **Bod1l** | 7.23E-10 | 0.56877783 | 0.473 | 0.109 | 1.49E-05 |
| **Nucks1** | 8.88E-10 | 0.69178037 | 0.538 | 0.153 | 1.83E-05 |
| **Mfsd14a** | 1.05E-09 | 0.46309292 | 0.264 | 0.022 | 2.16E-05 |
| **Ptprg** | 1.22E-09 | 0.62305267 | 0.341 | 0.055 | 2.51E-05 |
| **Prcp** | 1.34E-09 | 0.61962389 | 0.681 | 0.23 | 2.76E-05 |
| **Marveld1** | 1.58E-09 | 0.50556426 | 0.341 | 0.055 | 3.25E-05 |
| **Atp2b4** | 1.67E-09 | 0.61154847 | 0.473 | 0.12 | 3.43E-05 |
| **Ptges3** | 2.11E-09 | 0.75630568 | 0.604 | 0.202 | 4.34E-05 |
| **Zfp148** | 2.15E-09 | 0.35561182 | 0.407 | 0.082 | 4.44E-05 |
| **Gm42418** | 2.70E-09 | -1.287171 | 1 | 0.989 | 5.56E-05 |
| **Adgrl2** | 2.72E-09 | 0.42871249 | 0.352 | 0.06 | 5.61E-05 |
| **Tmem167** | 2.80E-09 | 0.34204668 | 0.505 | 0.131 | 5.77E-05 |
| **Pacs1** | 2.83E-09 | 0.53295284 | 0.253 | 0.022 | 5.84E-05 |
| **Dennd2a** | 3.18E-09 | 0.4592227 | 0.253 | 0.022 | 6.55E-05 |
| **Map1b** | 3.77E-09 | 0.4322794 | 0.308 | 0.044 | 7.77E-05 |
| **Emp2** | 3.90E-09 | 0.64222488 | 0.308 | 0.044 | 8.04E-05 |
| **Vim** | 4.24E-09 | 0.7117882 | 0.989 | 0.77 | 8.74E-05 |
| **Luzp1** | 4.85E-09 | 0.63469905 | 0.56 | 0.175 | 9.98E-05 |
| **Hif1a** | 4.87E-09 | 0.55471869 | 0.659 | 0.224 | 0.00010036 |
| **Lgmn** | 5.14E-09 | 0.55744764 | 0.418 | 0.093 | 0.00010588 |
| **Tax1bp3** | 5.27E-09 | 0.28949505 | 0.44 | 0.104 | 0.00010863 |
| **Mmp14** | 5.29E-09 | 0.40880163 | 0.484 | 0.126 | 0.00010889 |
| **Vegfc** | 5.93E-09 | 0.66120947 | 0.286 | 0.038 | 0.0001221 |
| **Mpp1** | 5.93E-09 | 0.58110912 | 0.385 | 0.082 | 0.0001222 |
| **Thsd1** | 6.86E-09 | 0.68723085 | 0.363 | 0.077 | 0.00014133 |
| **Irx3** | 7.09E-09 | 0.4148193 | 0.385 | 0.077 | 0.000146 |
| **Capg** | 7.86E-09 | 0.6012678 | 0.473 | 0.131 | 0.00016197 |
| **Dennd5b** | 7.92E-09 | 0.36531827 | 0.286 | 0.038 | 0.00016319 |
| **Ppic** | 8.86E-09 | 0.57696831 | 0.505 | 0.142 | 0.00018252 |
| **Nop56** | 1.01E-08 | 0.46753411 | 0.319 | 0.055 | 0.00020883 |
| **Ramp3** | 1.15E-08 | 0.68204812 | 0.33 | 0.06 | 0.00023618 |
| **Set** | 1.23E-08 | 0.49115651 | 0.659 | 0.24 | 0.00025395 |
| **Gatm** | 1.36E-08 | 0.66620684 | 0.429 | 0.109 | 0.00027999 |
| **Smim14** | 1.65E-08 | 0.56747796 | 0.462 | 0.126 | 0.00033933 |
| **Irf2bpl** | 1.74E-08 | 0.53268732 | 0.418 | 0.104 | 0.00035841 |
| **Pcp4l1** | 1.87E-08 | 0.62903124 | 0.78 | 0.328 | 0.00038597 |
| **Brd7** | 1.95E-08 | 0.32322402 | 0.44 | 0.109 | 0.00040207 |
| **Ube2r2** | 2.12E-08 | 0.41984037 | 0.549 | 0.164 | 0.00043653 |
| **Pi4k2a** | 2.12E-08 | 0.42464968 | 0.275 | 0.038 | 0.00043708 |
| **Svil** | 2.16E-08 | 0.3740386 | 0.615 | 0.202 | 0.00044445 |
| **Mapkap1** | 2.72E-08 | 0.43881404 | 0.286 | 0.044 | 0.00056102 |
| **Ythdf1** | 2.81E-08 | 0.34904869 | 0.319 | 0.055 | 0.0005799 |
| **Ube2v1** | 3.21E-08 | 0.43030945 | 0.396 | 0.093 | 0.00066134 |
| **She** | 3.69E-08 | 0.53628379 | 0.407 | 0.104 | 0.00076018 |
| **Dnaja2** | 3.78E-08 | 0.44251072 | 0.505 | 0.153 | 0.00077862 |
| **Xrn2** | 3.92E-08 | 0.41324887 | 0.429 | 0.115 | 0.00080806 |
| **Bag1** | 3.97E-08 | 0.44171391 | 0.484 | 0.148 | 0.00081762 |
| **Ap1s2** | 3.98E-08 | 0.38082783 | 0.308 | 0.055 | 0.00081939 |
| **Pbrm1** | 4.78E-08 | 0.43358129 | 0.516 | 0.153 | 0.00098388 |
| **Znrf1** | 5.31E-08 | 0.40791246 | 0.418 | 0.109 | 0.00109359 |
| **Rhoc** | 5.44E-08 | 0.69468811 | 0.857 | 0.426 | 0.00112007 |
| **Erh** | 5.56E-08 | 0.38881493 | 0.582 | 0.191 | 0.00114504 |
| **Bbx** | 5.80E-08 | 0.26194156 | 0.341 | 0.071 | 0.00119531 |
| **Dync1i2** | 6.29E-08 | 0.42449094 | 0.802 | 0.322 | 0.00129584 |
| **Rab14** | 6.43E-08 | 0.43759772 | 0.736 | 0.29 | 0.00132363 |
| **Fscn1** | 6.50E-08 | 0.42054733 | 0.527 | 0.175 | 0.00133901 |
| **Zfhx3** | 6.96E-08 | 0.4150661 | 0.516 | 0.164 | 0.00143386 |
| **Wipf1** | 7.19E-08 | 0.4860549 | 0.264 | 0.038 | 0.00148163 |
| **Manf** | 7.28E-08 | 0.36580161 | 0.593 | 0.202 | 0.00149915 |
| **Cmpk1** | 7.30E-08 | 0.37761663 | 0.462 | 0.131 | 0.00150302 |
| **Fgfr1** | 7.39E-08 | 0.38660076 | 0.451 | 0.126 | 0.00152302 |
| **Calm1** | 8.07E-08 | 0.59137033 | 0.989 | 0.874 | 0.00166295 |
| **Prrc2c** | 8.35E-08 | 0.362521 | 0.615 | 0.208 | 0.00172011 |
| **Ccdc50** | 8.76E-08 | 0.51399853 | 0.429 | 0.12 | 0.00180366 |
| **Rnaseh2c** | 8.81E-08 | 0.32240951 | 0.505 | 0.153 | 0.00181502 |
| **Pcmt1** | 8.97E-08 | 0.35470341 | 0.374 | 0.087 | 0.00184838 |
| **Arhgap5** | 9.02E-08 | 0.25268808 | 0.44 | 0.12 | 0.0018573 |
| **Timm17a** | 9.62E-08 | 0.48291 | 0.473 | 0.142 | 0.00198169 |
| **Caprin1** | 1.01E-07 | 0.3347238 | 0.516 | 0.153 | 0.00207406 |
| **Bmp2k** | 1.03E-07 | 0.53080245 | 0.363 | 0.087 | 0.0021238 |
| **Hdgf** | 1.04E-07 | 0.41666139 | 0.429 | 0.115 | 0.00213989 |
| **Anxa7** | 1.09E-07 | 0.56671983 | 0.505 | 0.153 | 0.00225406 |
| **Hecw2** | 1.27E-07 | 0.44108886 | 0.275 | 0.044 | 0.00261357 |
| **Rnf11** | 1.27E-07 | 0.45021502 | 0.396 | 0.104 | 0.00262338 |
| **Ltc4s** | 1.30E-07 | 0.33688474 | 0.527 | 0.18 | 0.00268483 |
| **Emid1** | 1.38E-07 | 0.50585195 | 0.451 | 0.131 | 0.0028441 |
| **Pdlim1** | 1.39E-07 | 0.65115459 | 0.538 | 0.191 | 0.002872 |
| **Ptpn11** | 1.40E-07 | 0.48993269 | 0.286 | 0.055 | 0.00288538 |
| **Gtf2a2** | 1.42E-07 | 0.48576487 | 0.505 | 0.164 | 0.00292101 |
| **mt-Cytb** | 1.43E-07 | -0.7890581 | 1 | 0.984 | 0.00295286 |
| **Odc1** | 1.48E-07 | 0.48008217 | 0.571 | 0.202 | 0.00304716 |
| **Vat1** | 1.51E-07 | 0.54735866 | 0.341 | 0.077 | 0.00310705 |
| **Ahnak2** | 1.51E-07 | 0.39956128 | 0.429 | 0.12 | 0.00311993 |
| **Peg13** | 1.52E-07 | 0.53836726 | 0.253 | 0.038 | 0.00313381 |
| **Metap2** | 1.53E-07 | 0.54863526 | 0.802 | 0.35 | 0.00315695 |
| **Pcm1** | 1.54E-07 | 0.3261868 | 0.484 | 0.142 | 0.00316903 |
| **Mlf2** | 1.59E-07 | 0.49321481 | 0.396 | 0.109 | 0.00327234 |
| **Adamts1** | 1.63E-07 | 0.7257233 | 0.538 | 0.197 | 0.0033521 |
| **Hnrnpa3** | 1.67E-07 | 0.45200564 | 0.736 | 0.301 | 0.00343301 |
| **Rbmxl1** | 1.73E-07 | 0.31651133 | 0.297 | 0.055 | 0.0035569 |
| **Ndel1** | 1.77E-07 | 0.27371706 | 0.374 | 0.093 | 0.00365198 |
| **Aebp1** | 1.79E-07 | 0.45362657 | 0.538 | 0.175 | 0.00368229 |
| **Anp32b** | 1.86E-07 | 0.48820918 | 0.549 | 0.191 | 0.00383054 |
| **Srp19** | 1.88E-07 | 0.31584533 | 0.429 | 0.12 | 0.00387398 |
| **Top2b** | 1.91E-07 | 0.34285309 | 0.418 | 0.115 | 0.00393495 |
| **Scoc** | 1.92E-07 | 0.47116703 | 0.44 | 0.126 | 0.00395737 |
| **Vps35** | 2.06E-07 | 0.43572692 | 0.429 | 0.126 | 0.00423599 |
| **Phlda3** | 2.12E-07 | 0.31941562 | 0.297 | 0.06 | 0.00436101 |
| **Palmd** | 2.12E-07 | 0.39231362 | 0.659 | 0.24 | 0.00436458 |
| **Anapc11** | 2.21E-07 | 0.32172538 | 0.505 | 0.164 | 0.0045501 |
| **Kras** | 2.32E-07 | 0.43296839 | 0.484 | 0.153 | 0.00477061 |
| **Ndufs4** | 2.34E-07 | 0.34671156 | 0.484 | 0.153 | 0.0048243 |
| **Kdelr1** | 2.36E-07 | 0.33349093 | 0.473 | 0.148 | 0.00485606 |
| **Taf1** | 2.39E-07 | 0.40562061 | 0.374 | 0.098 | 0.00492911 |
| **Spata6** | 2.44E-07 | 0.36951354 | 0.264 | 0.044 | 0.00503193 |
| **Ppp1r12a** | 2.44E-07 | 0.37746408 | 0.429 | 0.12 | 0.00503679 |
| **Pthlh** | 2.57E-07 | 0.70627703 | 0.275 | 0.049 | 0.00528816 |
| **Cdh13** | 2.59E-07 | 0.39799428 | 0.429 | 0.12 | 0.00534467 |
| **Fyn** | 2.65E-07 | 0.35791249 | 0.473 | 0.153 | 0.00546626 |
| **Pbdc1** | 2.68E-07 | 0.36909774 | 0.264 | 0.044 | 0.00552184 |
| **Pltp** | 2.73E-07 | 0.34451754 | 0.319 | 0.066 | 0.00561646 |
| **Uhrf2** | 2.79E-07 | 0.52491128 | 0.341 | 0.082 | 0.00574998 |
| **Sparc** | 2.80E-07 | 0.60531108 | 0.945 | 0.667 | 0.00576086 |
| **Dynlt3** | 2.95E-07 | 0.3377428 | 0.407 | 0.115 | 0.00607284 |
| **Slfn3** | 3.09E-07 | 0.54463332 | 0.308 | 0.066 | 0.00637473 |
| **Coro1c** | 3.14E-07 | 0.31211427 | 0.341 | 0.082 | 0.00647525 |
| **0610010K14Rik** | 3.16E-07 | 0.53878778 | 0.44 | 0.137 | 0.00650769 |
| **Cebpz** | 3.16E-07 | 0.46912362 | 0.462 | 0.142 | 0.00651251 |
| **Ldb2** | 3.19E-07 | 0.29878137 | 0.44 | 0.131 | 0.00657607 |
| **Hnrnpab** | 3.30E-07 | 0.33761627 | 0.571 | 0.202 | 0.006803 |
| **Cpne3** | 3.30E-07 | 0.34515575 | 0.297 | 0.06 | 0.0068056 |
| **Eif4e2** | 3.46E-07 | 0.38516508 | 0.308 | 0.066 | 0.00712697 |
| **Zfp560** | 3.46E-07 | 0.4556733 | 0.308 | 0.066 | 0.00712704 |
| **Rab9** | 3.54E-07 | 0.30727405 | 0.264 | 0.044 | 0.00728394 |
| **Golgb1** | 3.75E-07 | 0.53447961 | 0.462 | 0.148 | 0.00773078 |
| **Spred1** | 3.75E-07 | 0.42941874 | 0.297 | 0.06 | 0.00773447 |
| **Slfn2** | 3.83E-07 | 0.42639525 | 0.429 | 0.126 | 0.00789562 |
| **Sdhc** | 3.87E-07 | 0.35433583 | 0.308 | 0.066 | 0.00796471 |
| **Cd55** | 3.89E-07 | 0.62369379 | 0.264 | 0.049 | 0.00801331 |
| **Sox18** | 4.05E-07 | 0.474949 | 0.648 | 0.262 | 0.00833741 |
| **Bnip2** | 4.43E-07 | 0.47068157 | 0.516 | 0.186 | 0.00912124 |
| **Ppp1cb** | 4.43E-07 | 0.44220335 | 0.363 | 0.093 | 0.00912202 |
| **Dctn2** | 4.51E-07 | 0.40566857 | 0.538 | 0.186 | 0.00928492 |
| **Esf1** | 4.53E-07 | 0.26531082 | 0.407 | 0.115 | 0.00933523 |
| **Lrrfip1** | 4.63E-07 | 0.39774111 | 0.505 | 0.175 | 0.00954654 |
| **Ssna1** | 4.68E-07 | 0.32021101 | 0.385 | 0.104 | 0.00964308 |
| **Cd93** | 4.71E-07 | 0.32387772 | 0.473 | 0.148 | 0.00970399 |
| **Jcad** | 4.74E-07 | 0.52363471 | 0.462 | 0.153 | 0.00975766 |
| **Mff** | 4.84E-07 | 0.43162459 | 0.385 | 0.109 | 0.00996155 |
| **Rassf3** | 4.91E-07 | 0.46006029 | 0.385 | 0.104 | 0.01012394 |
| **Rock2** | 4.97E-07 | 0.37094602 | 0.549 | 0.202 | 0.01024697 |
| **Sf3b2** | 5.13E-07 | 0.41619366 | 0.549 | 0.202 | 0.0105658 |
| **Nr3c1** | 5.16E-07 | 0.38897484 | 0.549 | 0.191 | 0.01062748 |
| **Ccnd1** | 5.33E-07 | 0.53385275 | 0.418 | 0.131 | 0.01097547 |
| **Drg1** | 5.41E-07 | 0.40098341 | 0.286 | 0.06 | 0.01113638 |
| **Itgb4** | 5.43E-07 | 0.45231851 | 0.484 | 0.158 | 0.01119134 |
| **Lipa** | 5.46E-07 | 0.31028694 | 0.253 | 0.044 | 0.01123884 |
| **Smc3** | 5.68E-07 | 0.32629934 | 0.319 | 0.071 | 0.0116998 |
| **Wsb2** | 5.77E-07 | 0.49293276 | 0.319 | 0.077 | 0.01188241 |
| **Igfbp5** | 5.88E-07 | 0.84446196 | 0.857 | 0.459 | 0.01212044 |
| **Tm9sf2** | 6.20E-07 | 0.33449139 | 0.495 | 0.169 | 0.01276824 |
| **Tmed9** | 6.24E-07 | 0.43928239 | 0.571 | 0.219 | 0.01286395 |
| **Med13l** | 6.26E-07 | 0.41381373 | 0.253 | 0.044 | 0.01289284 |
| **Ahsa1** | 6.70E-07 | 0.46240556 | 0.352 | 0.093 | 0.01380477 |
| **Ttc28** | 6.78E-07 | 0.32404809 | 0.505 | 0.164 | 0.01397119 |
| **Clic5** | 6.80E-07 | 0.41992829 | 0.615 | 0.24 | 0.01401236 |
| **Psmd7** | 6.81E-07 | 0.48909657 | 0.484 | 0.164 | 0.01403612 |
| **Capn5** | 6.95E-07 | 0.37849025 | 0.275 | 0.055 | 0.01432421 |
| **Gm28875** | 7.05E-07 | 0.42826022 | 0.275 | 0.055 | 0.01453156 |
| **Uchl3** | 7.11E-07 | 0.36709032 | 0.505 | 0.169 | 0.01465388 |
| **Rbm25** | 7.21E-07 | 0.28700917 | 0.505 | 0.175 | 0.0148632 |
| **Mtmr6** | 7.38E-07 | 0.32289089 | 0.264 | 0.049 | 0.01519938 |
| **Wdr1** | 7.80E-07 | 0.47347023 | 0.33 | 0.082 | 0.01605889 |
| **Ube2k** | 7.80E-07 | 0.39376026 | 0.363 | 0.098 | 0.01606402 |
| **Tmed2** | 7.91E-07 | 0.40839989 | 0.659 | 0.273 | 0.01629912 |
| **Rab8b** | 7.93E-07 | 0.30624933 | 0.352 | 0.087 | 0.01633344 |
| **Pkd1** | 8.19E-07 | 0.27244379 | 0.363 | 0.098 | 0.01686859 |
| **Srp14** | 8.82E-07 | 0.43836307 | 0.692 | 0.29 | 0.01816291 |
| **Erbin** | 9.33E-07 | 0.29362072 | 0.33 | 0.082 | 0.01921769 |
| **Polr2c** | 9.38E-07 | 0.28901741 | 0.451 | 0.142 | 0.01931749 |
| **Rhoj** | 9.62E-07 | 0.49888421 | 0.637 | 0.262 | 0.01980954 |
| **Rnf130** | 1.00E-06 | 0.31168032 | 0.516 | 0.18 | 0.020611 |
| **Ccdc90b** | 1.00E-06 | 0.30473405 | 0.264 | 0.049 | 0.02066741 |
| **Mef2a** | 1.02E-06 | 0.36546068 | 0.648 | 0.251 | 0.02102831 |
| **Cd302** | 1.03E-06 | 0.3610102 | 0.484 | 0.164 | 0.02121535 |
| **Gps2** | 1.05E-06 | 0.33621007 | 0.352 | 0.093 | 0.02156844 |
| **Adam15** | 1.05E-06 | 0.26162618 | 0.516 | 0.18 | 0.02169303 |
| **Ptprk** | 1.07E-06 | 0.39497116 | 0.505 | 0.175 | 0.02205108 |
| **Ube2b** | 1.12E-06 | 0.58927264 | 0.593 | 0.246 | 0.02312023 |
| **Nfkbia** | 1.13E-06 | 0.65786405 | 0.879 | 0.53 | 0.02318669 |
| **Rcan1** | 1.13E-06 | 0.52308524 | 0.67 | 0.284 | 0.02321522 |
| **Ccny** | 1.13E-06 | 0.27851857 | 0.374 | 0.104 | 0.02329156 |
| **Midn** | 1.17E-06 | 0.28541419 | 0.352 | 0.093 | 0.0240861 |
| **Tfg** | 1.18E-06 | 0.35719074 | 0.308 | 0.071 | 0.02421637 |
| **Atp2b1** | 1.22E-06 | 0.39886627 | 0.538 | 0.197 | 0.02512288 |
| **Herpud1** | 1.23E-06 | 0.32006996 | 0.505 | 0.175 | 0.02533304 |
| **Thbd** | 1.24E-06 | 0.38743044 | 0.692 | 0.295 | 0.02547179 |
| **Ppp3ca** | 1.29E-06 | 0.28173625 | 0.527 | 0.186 | 0.026498 |
| **Pls3** | 1.39E-06 | 0.2995613 | 0.451 | 0.148 | 0.02863166 |
| **Cyfip1** | 1.41E-06 | 0.33009612 | 0.462 | 0.148 | 0.02896166 |
| **Ywhag** | 1.41E-06 | 0.37353508 | 0.253 | 0.049 | 0.02899962 |
| **Sf3b6** | 1.42E-06 | 0.29360066 | 0.538 | 0.197 | 0.02930398 |
| **Aamdc** | 1.43E-06 | 0.31153948 | 0.407 | 0.12 | 0.02948379 |
| **Rdx** | 1.43E-06 | 0.51716092 | 0.637 | 0.268 | 0.02955793 |
| **Sqstm1** | 1.44E-06 | 0.44311091 | 0.538 | 0.208 | 0.02964032 |
| **Grb2** | 1.47E-06 | 0.37466645 | 0.385 | 0.115 | 0.03020415 |
| **Gm16286** | 1.51E-06 | 0.37945275 | 0.484 | 0.169 | 0.03105271 |
| **Rraga** | 1.57E-06 | 0.3394401 | 0.363 | 0.104 | 0.0322406 |
| **Pum2** | 1.58E-06 | 0.46084954 | 0.396 | 0.126 | 0.03262993 |
| **Hsd17b11** | 1.65E-06 | 0.39197822 | 0.308 | 0.071 | 0.03403445 |
| **Nipbl** | 1.66E-06 | 0.45082716 | 0.396 | 0.126 | 0.03412958 |
| **Mcam** | 1.69E-06 | 0.29245353 | 0.297 | 0.066 | 0.03472027 |
| **Plat** | 1.69E-06 | 0.43341127 | 0.33 | 0.087 | 0.03475742 |
| **Zmynd8** | 1.70E-06 | 0.47831416 | 0.308 | 0.077 | 0.03501534 |
| **Uqcc2** | 1.71E-06 | 0.26130243 | 0.67 | 0.262 | 0.03519101 |
| **Pkig** | 1.75E-06 | 0.3705424 | 0.538 | 0.202 | 0.03598399 |
| **Arhgap29** | 1.75E-06 | 0.37269357 | 0.824 | 0.399 | 0.03601128 |
| **Bcas2** | 1.76E-06 | 0.31882866 | 0.44 | 0.142 | 0.03621659 |
| **Stap2** | 1.76E-06 | 0.38382457 | 0.297 | 0.071 | 0.03631202 |
| **Ppp5c** | 1.78E-06 | 0.33322327 | 0.253 | 0.049 | 0.03656894 |
| **Pgk1** | 1.80E-06 | 0.42416641 | 0.286 | 0.066 | 0.03697927 |
| **Lrrfip2** | 1.83E-06 | 0.25031362 | 0.253 | 0.049 | 0.03763907 |
| **Dnajc3** | 1.89E-06 | 0.34131304 | 0.549 | 0.202 | 0.03896832 |
| **S100a10** | 1.91E-06 | 0.57637492 | 0.978 | 0.77 | 0.0393912 |
| **Ctnnb1** | 1.97E-06 | 0.47256892 | 0.802 | 0.355 | 0.04053949 |
| **Zfp706** | 2.04E-06 | 0.43766969 | 0.571 | 0.23 | 0.04203842 |
| **Mesd** | 2.06E-06 | 0.41752252 | 0.308 | 0.077 | 0.04243421 |
| **Elf2** | 2.06E-06 | 0.33416916 | 0.538 | 0.202 | 0.04247493 |
| **Arl6ip5** | 2.07E-06 | 0.32443927 | 0.473 | 0.164 | 0.04271674 |
| **Glrx3** | 2.08E-06 | 0.3793749 | 0.44 | 0.148 | 0.04285766 |
| **Faim** | 2.14E-06 | 0.4315534 | 0.297 | 0.071 | 0.04416869 |
| **Pds5a** | 2.20E-06 | 0.27820421 | 0.308 | 0.077 | 0.04522678 |
| **Mrpl21** | 2.21E-06 | 0.40093241 | 0.319 | 0.082 | 0.04548552 |
| **Clint1** | 2.27E-06 | 0.44390744 | 0.264 | 0.055 | 0.04685898 |
| **Naxd** | 2.31E-06 | 0.36446467 | 0.286 | 0.066 | 0.04762996 |
| **Rtl8a** | 2.32E-06 | 0.31864434 | 0.527 | 0.197 | 0.04769673 |
| **Tacc1** | 2.32E-06 | 0.39566063 | 0.538 | 0.202 | 0.04771891 |
| **Sema3f** | 2.33E-06 | 0.43006017 | 0.604 | 0.257 | 0.04798278 |
| **Ankrd40** | 2.34E-06 | 0.29333937 | 0.264 | 0.055 | 0.04818069 |
| **Sap30l** | 2.34E-06 | 0.28575963 | 0.308 | 0.077 | 0.04819538 |
| **Anapc16** | 2.35E-06 | 0.44656399 | 0.484 | 0.18 | 0.04850483 |
| **Polb** | 2.40E-06 | 0.28796869 | 0.253 | 0.049 | 0.04942381 |
| **Smarca5** | 2.42E-06 | 0.40497661 | 0.429 | 0.142 | 0.04995715 |
| **Hspa1a** | 2.43E-06 | 0.79490142 | 0.538 | 0.219 | 0.04997759 |
| **Nrip1** | 2.43E-06 | 0.27179427 | 0.407 | 0.126 | 0.05014776 |
| **Txndc15** | 2.48E-06 | 0.27690215 | 0.341 | 0.093 | 0.05106349 |
| **Daam1** | 2.51E-06 | 0.38137425 | 0.264 | 0.055 | 0.05164341 |
| **Pitpnb** | 2.53E-06 | 0.2812545 | 0.319 | 0.082 | 0.05216098 |
| **Tcaf1** | 2.54E-06 | 0.31391149 | 0.297 | 0.071 | 0.05227611 |
| **Nrbp1** | 2.63E-06 | 0.3390037 | 0.407 | 0.126 | 0.05414919 |
| **Yy1** | 2.64E-06 | 0.43663665 | 0.418 | 0.137 | 0.05429707 |
| **Spop** | 2.65E-06 | 0.31242103 | 0.473 | 0.164 | 0.05460567 |
| **Agpat4** | 2.70E-06 | 0.30781283 | 0.352 | 0.098 | 0.05555379 |
| **Col4a2** | 2.72E-06 | 0.30773113 | 0.659 | 0.279 | 0.05594876 |
| **Smarcb1** | 2.73E-06 | 0.31159431 | 0.275 | 0.06 | 0.05621124 |
| **Chd1** | 2.78E-06 | 0.31378798 | 0.297 | 0.071 | 0.05721566 |
| **Tiprl** | 2.79E-06 | 0.25822147 | 0.308 | 0.077 | 0.05753624 |
| **Psmb4** | 2.84E-06 | 0.31159398 | 0.549 | 0.202 | 0.0584233 |
| **Kdelr2** | 2.85E-06 | 0.39510663 | 0.396 | 0.126 | 0.05880723 |
| **Vdac2** | 2.90E-06 | 0.31715359 | 0.593 | 0.224 | 0.05978519 |
| **Dpysl2** | 2.91E-06 | 0.34084575 | 0.385 | 0.115 | 0.06003114 |
| **Rab12** | 2.94E-06 | 0.34112627 | 0.407 | 0.131 | 0.06054886 |
| **Cdv3** | 3.03E-06 | 0.35663927 | 0.429 | 0.142 | 0.06242679 |
| **Mbtps1** | 3.04E-06 | 0.27194548 | 0.275 | 0.06 | 0.0625955 |
| **Tgfb1** | 3.07E-06 | 0.30193333 | 0.484 | 0.175 | 0.06328461 |
| **Ano6** | 3.17E-06 | 0.26219221 | 0.264 | 0.055 | 0.06529382 |
| **Bin1** | 3.21E-06 | 0.28423772 | 0.538 | 0.202 | 0.06617146 |
| **Mindy2** | 3.22E-06 | 0.4181635 | 0.396 | 0.126 | 0.06634027 |
| **Clic4** | 3.31E-06 | 0.50817161 | 0.769 | 0.366 | 0.06813913 |
| **Kctd12** | 3.35E-06 | 0.52643451 | 0.527 | 0.208 | 0.06907516 |
| **Pbx1** | 3.53E-06 | 0.28063827 | 0.319 | 0.082 | 0.07276903 |
| **Foxp1** | 3.55E-06 | 0.37340128 | 0.681 | 0.284 | 0.07318803 |
| **Cxcl1** | 3.66E-06 | 0.86074239 | 0.396 | 0.131 | 0.0754507 |
| **Hectd1** | 3.68E-06 | 0.486044 | 0.374 | 0.12 | 0.07586388 |
| **Ccser2** | 3.72E-06 | 0.26540382 | 0.462 | 0.164 | 0.07669594 |
| **Rab11b** | 3.83E-06 | 0.26915996 | 0.44 | 0.153 | 0.07887739 |
| **Pgrmc1** | 3.87E-06 | 0.32232612 | 0.451 | 0.153 | 0.07973776 |
| **Plec** | 3.88E-06 | 0.51949348 | 0.802 | 0.41 | 0.07987242 |
| **Apopt1** | 3.90E-06 | 0.28580094 | 0.429 | 0.142 | 0.08033453 |
| **Aip** | 3.94E-06 | 0.32373152 | 0.297 | 0.077 | 0.08116803 |
| **Sec11a** | 4.02E-06 | 0.33073333 | 0.484 | 0.175 | 0.08272935 |
| **Pkn1** | 4.09E-06 | 0.34966728 | 0.297 | 0.077 | 0.08428271 |
| **Ndufb2** | 4.12E-06 | 0.29165187 | 0.549 | 0.208 | 0.08477397 |
| **Tial1** | 4.20E-06 | 0.40858528 | 0.33 | 0.093 | 0.08644068 |
| **Akt1** | 4.44E-06 | 0.40149211 | 0.308 | 0.082 | 0.09143575 |
| **Map3k8** | 4.50E-06 | 0.31160995 | 0.33 | 0.093 | 0.09279145 |
| **Hnrnpa0** | 4.51E-06 | 0.40611964 | 0.637 | 0.262 | 0.0928311 |
| **Arpc5l** | 4.52E-06 | 0.27586389 | 0.407 | 0.131 | 0.09303115 |
| **Oser1** | 4.52E-06 | 0.25529059 | 0.297 | 0.077 | 0.09315646 |
| **Larp4** | 4.67E-06 | 0.41789847 | 0.33 | 0.093 | 0.09613321 |
| **Wdr26** | 4.68E-06 | 0.38181074 | 0.363 | 0.109 | 0.09642281 |
| **Rb1cc1** | 4.71E-06 | 0.28004664 | 0.341 | 0.098 | 0.09704041 |
| **Itch** | 4.75E-06 | 0.27432088 | 0.275 | 0.066 | 0.09795738 |
| **Glrx5** | 4.89E-06 | 0.28785583 | 0.33 | 0.093 | 0.10076759 |
| **Chmp3** | 4.94E-06 | 0.30154441 | 0.593 | 0.235 | 0.10170453 |
| **Bvht** | 4.95E-06 | 0.31871848 | 0.56 | 0.219 | 0.10192387 |
| **Mib1** | 4.95E-06 | 0.30878471 | 0.286 | 0.071 | 0.1020436 |
| **Btbd3** | 4.99E-06 | 0.35807576 | 0.582 | 0.23 | 0.10281713 |
| **Ube2d3** | 5.31E-06 | 0.4803212 | 0.681 | 0.306 | 0.10933465 |
| **Pgam1** | 5.41E-06 | 0.28559694 | 0.44 | 0.158 | 0.1114948 |
| **Ddx23** | 5.41E-06 | 0.32375921 | 0.275 | 0.066 | 0.11153551 |
| **Arih1** | 5.52E-06 | 0.26289681 | 0.297 | 0.077 | 0.11366218 |
| **Mindy1** | 5.53E-06 | 0.30600049 | 0.308 | 0.082 | 0.11387549 |
| **Synpo** | 5.58E-06 | 0.35055156 | 0.44 | 0.158 | 0.11491809 |
| **Hacd4** | 5.58E-06 | 0.2852708 | 0.253 | 0.055 | 0.11498713 |
| **Huwe1** | 5.63E-06 | 0.34337127 | 0.385 | 0.126 | 0.11603422 |
| **Sh3bgrl** | 5.70E-06 | 0.46648599 | 0.429 | 0.153 | 0.11740153 |
| **Nfyb** | 5.81E-06 | 0.35096323 | 0.253 | 0.055 | 0.1197663 |
| **Irf2** | 5.84E-06 | 0.30096058 | 0.462 | 0.164 | 0.12026064 |
| **Ruvbl1** | 5.97E-06 | 0.2725941 | 0.253 | 0.055 | 0.12305742 |
| **Evi5** | 6.00E-06 | 0.26920088 | 0.341 | 0.098 | 0.12359951 |
| **Supt16** | 6.07E-06 | 0.33812407 | 0.363 | 0.115 | 0.12508762 |
| **Lmo4** | 6.16E-06 | 0.44557435 | 0.308 | 0.082 | 0.1269859 |
| **Tob1** | 6.24E-06 | 0.25725835 | 0.418 | 0.142 | 0.12864516 |
| **Ppdpf** | 6.28E-06 | 0.39860883 | 0.538 | 0.213 | 0.12943496 |
| **Sod1** | 6.28E-06 | 0.27241634 | 0.56 | 0.224 | 0.12943953 |
| **Kxd1** | 6.30E-06 | 0.29665847 | 0.352 | 0.104 | 0.1298551 |
| **St6galnac4** | 6.38E-06 | 0.25374553 | 0.286 | 0.071 | 0.13138574 |
| **Creg1** | 6.40E-06 | 0.41503942 | 0.297 | 0.077 | 0.13180767 |
| **Homer3** | 6.40E-06 | 0.33766252 | 0.33 | 0.093 | 0.13185775 |
| **Zbtb7c** | 6.65E-06 | 0.25024664 | 0.275 | 0.066 | 0.13707858 |
| **Srebf2** | 6.71E-06 | 0.29092281 | 0.308 | 0.082 | 0.13816917 |
| **Nhp2** | 6.85E-06 | 0.43675145 | 0.363 | 0.115 | 0.14115913 |
| **Gmfb** | 6.89E-06 | 0.29284594 | 0.297 | 0.077 | 0.1418893 |
| **Morf4l1** | 6.95E-06 | 0.45734565 | 0.879 | 0.448 | 0.14315028 |
| **Sumo1** | 7.02E-06 | 0.34883115 | 0.593 | 0.257 | 0.14470929 |
| **Tpr** | 7.05E-06 | 0.29361333 | 0.615 | 0.251 | 0.14528362 |
| **Ctnnd1** | 7.12E-06 | 0.33205083 | 0.462 | 0.169 | 0.14659307 |
| **Ccdc12** | 7.16E-06 | 0.36823277 | 0.582 | 0.24 | 0.14752208 |
| **Srpk2** | 7.28E-06 | 0.32801398 | 0.385 | 0.126 | 0.14996974 |
| **Arl13b** | 7.33E-06 | 0.27417175 | 0.319 | 0.087 | 0.15107943 |
| **Rnf44** | 7.46E-06 | 0.45110181 | 0.275 | 0.071 | 0.15360683 |
| **Hspe1** | 7.51E-06 | 0.2602288 | 0.495 | 0.18 | 0.1546773 |
| **Cd34** | 7.52E-06 | 0.48892561 | 0.912 | 0.508 | 0.15500524 |
| **Smarce1** | 7.54E-06 | 0.31044234 | 0.418 | 0.148 | 0.15539967 |
| **Tspan13** | 7.55E-06 | 0.26060086 | 0.44 | 0.153 | 0.15562505 |
| **Ptpn1** | 7.56E-06 | 0.26505159 | 0.505 | 0.186 | 0.15568311 |
| **Sec63** | 7.79E-06 | 0.35043817 | 0.396 | 0.131 | 0.16051089 |
| **Mnat1** | 7.89E-06 | 0.255411 | 0.286 | 0.071 | 0.16252275 |
| **Selenof** | 7.97E-06 | 0.57398897 | 0.703 | 0.328 | 0.16415322 |
| **Phf11d** | 8.26E-06 | 0.30444149 | 0.319 | 0.093 | 0.17024695 |
| **Entpd1** | 8.27E-06 | 0.30324204 | 0.297 | 0.077 | 0.17042161 |
| **Sppl2a** | 8.30E-06 | 0.27711873 | 0.374 | 0.12 | 0.17093014 |
| **Eif1ax** | 8.32E-06 | 0.27236576 | 0.462 | 0.169 | 0.17130203 |
| **Anp32a** | 8.36E-06 | 0.37932083 | 0.681 | 0.29 | 0.17231791 |
| **mt-Co2** | 8.85E-06 | -0.6643316 | 1 | 0.995 | 0.18237524 |
| **Eef1g** | 8.86E-06 | 0.37767795 | 0.571 | 0.235 | 0.18260624 |
| **mt-Co3** | 8.92E-06 | -0.5898435 | 1 | 1 | 0.18375129 |
| **Fndc3b** | 9.19E-06 | 0.33523133 | 0.363 | 0.115 | 0.18941136 |
| **Cnot6** | 9.39E-06 | 0.26252896 | 0.308 | 0.087 | 0.19345236 |
| **Lsm6** | 9.41E-06 | 0.32943574 | 0.473 | 0.169 | 0.19386291 |
| **Polr2m** | 9.55E-06 | 0.25035487 | 0.407 | 0.142 | 0.19663748 |
| **Rassf1** | 9.90E-06 | 0.28939965 | 0.462 | 0.164 | 0.20401751 |
| **Eea1** | 9.92E-06 | 0.29359144 | 0.363 | 0.115 | 0.20428773 |
| **Gas6** | 1.01E-05 | 0.29129848 | 0.703 | 0.311 | 0.20749214 |
| **Phc2** | 1.09E-05 | 0.33223455 | 0.363 | 0.115 | 0.22505947 |
| **Nudt3** | 1.10E-05 | 0.50710246 | 0.264 | 0.066 | 0.22652524 |
| **Atrx** | 1.10E-05 | 0.25551952 | 0.484 | 0.186 | 0.22672659 |
| **Vgll4** | 1.11E-05 | 0.31673093 | 0.516 | 0.197 | 0.22792671 |
| **Cux1** | 1.11E-05 | 0.3459944 | 0.341 | 0.109 | 0.22888798 |
| **Wasl** | 1.12E-05 | 0.2732498 | 0.418 | 0.142 | 0.22978875 |
| **Lst1** | 1.12E-05 | 0.37120635 | 0.253 | 0.06 | 0.23093362 |
| **Endod1** | 1.14E-05 | 0.28487326 | 0.396 | 0.131 | 0.23560124 |
| **Tnfrsf1a** | 1.17E-05 | 0.39432531 | 0.516 | 0.202 | 0.24034311 |
| **Nptn** | 1.17E-05 | 0.31787723 | 0.33 | 0.098 | 0.24176392 |
| **Ednrb** | 1.18E-05 | 0.44737652 | 0.505 | 0.202 | 0.2430057 |
| **Fbxl7** | 1.20E-05 | 0.27740792 | 0.253 | 0.06 | 0.2464252 |
| **Cast** | 1.25E-05 | 0.26667314 | 0.593 | 0.246 | 0.25701218 |
| **Zfp771** | 1.26E-05 | 0.29660863 | 0.264 | 0.066 | 0.26036621 |
| **Bcr** | 1.28E-05 | 0.57493932 | 0.56 | 0.23 | 0.26403609 |
| **Ywhah** | 1.28E-05 | 0.31806244 | 0.604 | 0.262 | 0.26420474 |
| **Bcap31** | 1.31E-05 | 0.26665259 | 0.495 | 0.186 | 0.27042479 |
| **Ran** | 1.32E-05 | 0.29134382 | 0.714 | 0.317 | 0.27284434 |
| **Pnpla2** | 1.33E-05 | 0.29693805 | 0.253 | 0.06 | 0.27329316 |
| **Zscan26** | 1.33E-05 | 0.31410714 | 0.253 | 0.06 | 0.27329316 |
| **Rpl36a** | 1.34E-05 | 0.52154145 | 0.923 | 0.59 | 0.27528874 |
| **Yipf3** | 1.38E-05 | 0.28434192 | 0.352 | 0.115 | 0.28387329 |
| **Ppm1a** | 1.38E-05 | 0.25711494 | 0.264 | 0.066 | 0.28435752 |
| **Pde4b** | 1.41E-05 | 0.3566075 | 0.374 | 0.126 | 0.29049421 |
| **Helz** | 1.44E-05 | 0.25044009 | 0.297 | 0.082 | 0.29597047 |
| **Arpc1b** | 1.46E-05 | 0.46539515 | 0.824 | 0.415 | 0.30146462 |
| **Med19** | 1.51E-05 | 0.2743982 | 0.286 | 0.077 | 0.31162538 |
| **Stx16** | 1.54E-05 | 0.33797785 | 0.264 | 0.066 | 0.31632643 |
| **Dusp6** | 1.57E-05 | 0.29596834 | 0.429 | 0.153 | 0.32385684 |
| **Atg3** | 1.57E-05 | 0.29615593 | 0.33 | 0.104 | 0.32434463 |
| **Gdi2** | 1.64E-05 | 0.32467384 | 0.593 | 0.262 | 0.33746484 |
| **Fxyd5** | 1.69E-05 | 0.39594576 | 0.791 | 0.404 | 0.34739299 |
| **Pold4** | 1.69E-05 | 0.31108845 | 0.33 | 0.098 | 0.34900421 |
| **Sema3a** | 1.87E-05 | 0.36912726 | 0.516 | 0.208 | 0.38552326 |
| **Tbrg1** | 1.88E-05 | 0.27704574 | 0.495 | 0.197 | 0.38691553 |
| **mt-Co1** | 1.88E-05 | -0.5427424 | 1 | 0.995 | 0.38790146 |
| **Timm13** | 1.93E-05 | 0.29500529 | 0.538 | 0.219 | 0.39803136 |
| **Flrt2** | 1.94E-05 | 0.29286035 | 0.418 | 0.148 | 0.39989489 |
| **Psmc3** | 1.95E-05 | 0.26340548 | 0.484 | 0.191 | 0.40099679 |
| **Xbp1** | 1.97E-05 | 0.36431839 | 0.319 | 0.098 | 0.40502944 |
| **Cript** | 2.05E-05 | 0.35255398 | 0.451 | 0.175 | 0.42258483 |
| **Epas1** | 2.07E-05 | 0.26383451 | 0.571 | 0.24 | 0.42686266 |
| **Ppp1r16b** | 2.08E-05 | 0.25470282 | 0.341 | 0.109 | 0.42863236 |
| **Bcl10** | 2.13E-05 | 0.50133885 | 0.429 | 0.164 | 0.43866205 |
| **1110038F14Rik** | 2.19E-05 | 0.25452477 | 0.418 | 0.153 | 0.45197752 |
| **Zfp207** | 2.36E-05 | 0.26616879 | 0.341 | 0.109 | 0.48692582 |
| **Nfe2l1** | 2.47E-05 | 0.36914711 | 0.736 | 0.35 | 0.50946564 |
| **Trpc4ap** | 2.51E-05 | 0.25316727 | 0.297 | 0.087 | 0.5171996 |
| **Ppid** | 2.51E-05 | 0.29115133 | 0.297 | 0.087 | 0.5171996 |
| **Mrpl43** | 2.58E-05 | 0.29431615 | 0.352 | 0.115 | 0.53162088 |
| **Ywhaz** | 2.65E-05 | 0.36782762 | 0.681 | 0.322 | 0.54532754 |
| **Nme1** | 2.67E-05 | 0.26263484 | 0.549 | 0.235 | 0.54958424 |
| **Anxa3** | 2.67E-05 | 0.42959105 | 0.67 | 0.322 | 0.55002322 |
| **Nupr1** | 2.74E-05 | 0.51677544 | 0.934 | 0.705 | 0.56366593 |
| **Ntn1** | 2.74E-05 | 0.27377062 | 0.648 | 0.284 | 0.5649773 |
| **Dlc1** | 2.76E-05 | 0.27922684 | 0.33 | 0.104 | 0.56960752 |
| **Ssb** | 2.87E-05 | 0.30019955 | 0.56 | 0.251 | 0.59029636 |
| **Ptbp1** | 3.00E-05 | 0.2904426 | 0.341 | 0.115 | 0.6181413 |
| **Hikeshi** | 3.08E-05 | 0.31341929 | 0.352 | 0.115 | 0.63381347 |
| **Rad23a** | 3.12E-05 | 0.25020149 | 0.275 | 0.077 | 0.64301447 |
| **Akap13** | 3.16E-05 | 0.34074159 | 0.703 | 0.328 | 0.65129081 |
| **Tpm4** | 3.20E-05 | 0.48144289 | 0.769 | 0.443 | 0.66005945 |
| **F2r** | 3.30E-05 | 0.35597725 | 0.473 | 0.191 | 0.68066421 |
| **Tmem128** | 3.36E-05 | 0.30378914 | 0.286 | 0.082 | 0.6921783 |
| **Cul1** | 3.43E-05 | 0.31267454 | 0.275 | 0.077 | 0.7058237 |
| **Ncor2** | 3.45E-05 | 0.35550339 | 0.319 | 0.104 | 0.71002692 |
| **Mxra8** | 3.52E-05 | 0.259159 | 0.747 | 0.344 | 0.72535874 |
| **Guk1** | 3.53E-05 | 0.25663911 | 0.429 | 0.164 | 0.72736484 |
| **Wdr45b** | 3.78E-05 | 0.29582455 | 0.33 | 0.109 | 0.77829977 |
| **Ifnar1** | 3.84E-05 | 0.33834974 | 0.253 | 0.066 | 0.79091774 |
| **Lamb1** | 3.87E-05 | 0.32970136 | 0.319 | 0.104 | 0.79805172 |
| **Samsn1** | 3.99E-05 | 0.47671488 | 0.418 | 0.164 | 0.82166269 |
| **Tecr** | 3.99E-05 | 0.25577649 | 0.418 | 0.158 | 0.82216135 |
| **Upf3b** | 4.00E-05 | 0.31337481 | 0.319 | 0.104 | 0.82379967 |
| **Med15** | 4.03E-05 | 0.30365173 | 0.253 | 0.071 | 0.83040448 |
| **Cdc42** | 4.07E-05 | 0.36176197 | 0.802 | 0.393 | 0.83886113 |
| **Cebpzos** | 4.11E-05 | 0.27229532 | 0.264 | 0.071 | 0.84686302 |
| **Ypel2** | 4.15E-05 | 0.25938065 | 0.308 | 0.098 | 0.85506071 |
| **Mfsd4a** | 4.16E-05 | 0.25564291 | 0.352 | 0.12 | 0.85659587 |
| **Akirin1** | 4.22E-05 | 0.28394726 | 0.341 | 0.115 | 0.86839225 |
| **Tmem258** | 4.46E-05 | 0.25567466 | 0.473 | 0.186 | 0.91911175 |
| **Txnl1** | 4.74E-05 | 0.27861349 | 0.527 | 0.23 | 0.97546089 |
| **Cct4** | 4.83E-05 | 0.32103235 | 0.44 | 0.18 | 0.99527434 |
| **Rheb** | 4.87E-05 | 0.36524693 | 0.67 | 0.311 | 1 |
| **Msn** | 5.19E-05 | 0.40942448 | 0.813 | 0.41 | 1 |
| **Maf** | 5.19E-05 | 0.33860655 | 0.692 | 0.35 | 1 |
| **Hnrnpd** | 5.49E-05 | 0.27648366 | 0.473 | 0.191 | 1 |
| **Atp6v1d** | 5.77E-05 | 0.25913675 | 0.319 | 0.104 | 1 |
| **Slc25a3** | 5.80E-05 | 0.36128405 | 0.813 | 0.399 | 1 |
| **Ehd2** | 5.80E-05 | 0.35867996 | 0.626 | 0.284 | 1 |
| **Col4a1** | 5.93E-05 | 0.45631886 | 0.714 | 0.355 | 1 |
| **Etfa** | 6.44E-05 | 0.32877519 | 0.341 | 0.126 | 1 |
| **Samd4b** | 6.58E-05 | 0.33283949 | 0.319 | 0.109 | 1 |
| **Phldb2** | 6.74E-05 | 0.33574696 | 0.396 | 0.153 | 1 |
| **Sugt1** | 6.74E-05 | 0.37758546 | 0.538 | 0.24 | 1 |
| **Itgb1** | 6.95E-05 | 0.51828203 | 0.857 | 0.536 | 1 |
| **Sec24b** | 6.96E-05 | 0.28191702 | 0.264 | 0.077 | 1 |
| **Flna** | 7.12E-05 | 0.31809714 | 0.33 | 0.115 | 1 |
| **Timm23** | 7.12E-05 | 0.32832918 | 0.308 | 0.098 | 1 |
| **Eif5b** | 7.25E-05 | 0.2624094 | 0.549 | 0.23 | 1 |
| **Slc30a4** | 7.25E-05 | 0.27602293 | 0.253 | 0.071 | 1 |
| **Taok1** | 7.27E-05 | 0.27205872 | 0.374 | 0.137 | 1 |
| **Hivep2** | 7.59E-05 | 0.31821209 | 0.319 | 0.109 | 1 |
| **Atg12** | 8.09E-05 | 0.29716993 | 0.286 | 0.087 | 1 |
| **Arf5** | 8.81E-05 | 0.37885076 | 0.571 | 0.273 | 1 |
| **Jak1** | 9.01E-05 | 0.30584343 | 0.571 | 0.262 | 1 |
| **Marcksl1** | 9.30E-05 | 0.25679407 | 0.582 | 0.262 | 1 |
| **Papola** | 9.78E-05 | 0.27327644 | 0.33 | 0.115 | 1 |
| **Ube2h** | 0.00010606 | 0.32216719 | 0.275 | 0.087 | 1 |
| **Stk16** | 0.00010942 | 0.36531038 | 0.264 | 0.082 | 1 |
| **Ssbp4** | 0.00011101 | 0.25243114 | 0.396 | 0.153 | 1 |
| **Rbm22** | 0.0001118 | 0.27704779 | 0.352 | 0.131 | 1 |
| **Ywhaq** | 0.00011404 | 0.2859448 | 0.725 | 0.339 | 1 |
| **Qk** | 0.0001142 | 0.32057858 | 0.659 | 0.322 | 1 |
| **M6pr** | 0.00011431 | 0.29315051 | 0.264 | 0.082 | 1 |
| **Sh3bp5** | 0.00011432 | 0.3558861 | 0.516 | 0.235 | 1 |
| **Pecam1** | 0.00011442 | 0.32693162 | 0.835 | 0.448 | 1 |
| **Dag1** | 0.00011553 | 0.31543314 | 0.275 | 0.087 | 1 |
| **Wipi1** | 0.00011676 | 0.261072 | 0.275 | 0.087 | 1 |
| **Dst** | 0.00012035 | 0.27583572 | 0.703 | 0.35 | 1 |
| **Gm13889** | 0.00012189 | 0.63890704 | 0.253 | 0.077 | 1 |
| **Rpl32** | 0.00012469 | 0.37137534 | 0.978 | 0.792 | 1 |
| **Cd81** | 0.00012558 | 0.48334488 | 0.802 | 0.492 | 1 |
| **Vapa** | 0.00012758 | 0.25256404 | 0.56 | 0.257 | 1 |
| **Ifrd1** | 0.00012891 | 0.27216132 | 0.484 | 0.208 | 1 |
| **Hmcn1** | 0.00012919 | 0.29378555 | 0.879 | 0.503 | 1 |
| **Pdia6** | 0.00012948 | 0.2836628 | 0.495 | 0.219 | 1 |
| **Rpl39** | 0.00013039 | 0.34420715 | 1 | 0.896 | 1 |
| **Ifi27l2a** | 0.00013176 | 0.45785598 | 0.78 | 0.47 | 1 |
| **Gas5** | 0.00014163 | 0.28336748 | 0.615 | 0.284 | 1 |
| **Crip1** | 0.00014606 | 0.64889437 | 0.692 | 0.388 | 1 |
| **Ctla2a** | 0.0001574 | 0.41692994 | 0.846 | 0.497 | 1 |
| **Rere** | 0.00015898 | 0.2785797 | 0.286 | 0.093 | 1 |
| **Trim44** | 0.00016409 | 0.28232612 | 0.253 | 0.077 | 1 |
| **mt-Nd4** | 0.00016587 | -0.5777086 | 1 | 0.967 | 1 |
| **Avpi1** | 0.00017456 | 0.26445996 | 0.418 | 0.18 | 1 |
| **Camk1** | 0.00017907 | 0.29341351 | 0.275 | 0.093 | 1 |
| **Ptgds** | 0.00018437 | -1.5089853 | 0.505 | 0.596 | 1 |
| **Purb** | 0.00020118 | 0.25700436 | 0.462 | 0.208 | 1 |
| **Kdm1a** | 0.00020124 | 0.28084226 | 0.341 | 0.126 | 1 |
| **Cav1** | 0.00020914 | 0.28599385 | 0.615 | 0.311 | 1 |
| **Yaf2** | 0.00021444 | 0.27117738 | 0.286 | 0.098 | 1 |
| **Ppp4r2** | 0.00022331 | 0.25167702 | 0.33 | 0.126 | 1 |
| **Rab11a** | 0.00022681 | 0.35294819 | 0.813 | 0.437 | 1 |
| **Brd2** | 0.00022815 | 0.26704486 | 0.473 | 0.213 | 1 |
| **Pnp** | 0.00023625 | 0.30051971 | 0.462 | 0.197 | 1 |
| **Rpl22l1** | 0.00023771 | 0.39927681 | 0.956 | 0.552 | 1 |
| **Fkbp1a** | 0.00026587 | 0.37891822 | 0.934 | 0.601 | 1 |
| **Sptan1** | 0.00028761 | 0.26071186 | 0.615 | 0.295 | 1 |
| **Arhgdib** | 0.00028843 | 0.27103096 | 0.275 | 0.093 | 1 |
| **H2-D1** | 0.00029569 | 0.42993264 | 0.934 | 0.65 | 1 |
| **Tuba1a** | 0.00030095 | 0.36012217 | 0.495 | 0.235 | 1 |
| **Sf3a3** | 0.00030325 | 0.26179684 | 0.264 | 0.087 | 1 |
| **Sars** | 0.00030914 | 0.27662819 | 0.407 | 0.175 | 1 |
| **AA467197** | 0.00030928 | 0.366702 | 0.275 | 0.093 | 1 |
| **Pon2** | 0.00031264 | 0.25437999 | 0.264 | 0.087 | 1 |
| **Hnrnpk** | 0.00031831 | 0.27774009 | 0.747 | 0.383 | 1 |
| **Sfr1** | 0.0003615 | 0.25848065 | 0.659 | 0.322 | 1 |
| **Slc9a3r2** | 0.0003688 | 0.34316578 | 0.846 | 0.497 | 1 |
| **Hspa1b** | 0.00037883 | 0.34319231 | 0.648 | 0.339 | 1 |
| **Tmem30a** | 0.00038147 | 0.25735218 | 0.549 | 0.262 | 1 |
| **mt-Nd1** | 0.00039986 | -0.6456462 | 0.978 | 0.896 | 1 |
| **mt-Atp6** | 0.00040726 | -0.5674275 | 1 | 1 | 1 |
| **Tspan31** | 0.00042068 | 0.25860429 | 0.264 | 0.093 | 1 |
| **Cfl1** | 0.00044339 | 0.31060529 | 0.791 | 0.421 | 1 |
| **Rpl15** | 0.0004542 | 0.31316693 | 0.967 | 0.765 | 1 |
| **Clic1** | 0.00049646 | 0.3394234 | 0.934 | 0.607 | 1 |
| **Hsp90ab1** | 0.00051576 | 0.31986037 | 0.978 | 0.809 | 1 |
| **Cdk11b** | 0.00054782 | 0.29891281 | 0.429 | 0.197 | 1 |
| **Serpine1** | 0.00056717 | 0.38879204 | 0.275 | 0.104 | 1 |
| **Crk** | 0.00056994 | 0.28889166 | 0.341 | 0.142 | 1 |
| **Mrfap1** | 0.00059152 | 0.26937871 | 0.549 | 0.273 | 1 |
| **Rps18** | 0.00062628 | 0.29917659 | 0.945 | 0.743 | 1 |
| **Rac1** | 0.00066011 | 0.29445536 | 0.725 | 0.388 | 1 |
| **Cox5a** | 0.00068054 | 0.30390342 | 0.703 | 0.377 | 1 |
| **Rps12** | 0.00070064 | 0.35536928 | 0.923 | 0.585 | 1 |
| **Reep3** | 0.00079646 | 0.32948092 | 0.648 | 0.339 | 1 |
| **Rpsa** | 0.000818 | 0.32149881 | 0.956 | 0.727 | 1 |
| **Adamtsl1** | 0.0008224 | 0.26313707 | 0.89 | 0.497 | 1 |
| **Fxyd6** | 0.00083291 | -0.5522388 | 0.813 | 0.77 | 1 |
| **Pafah1b2** | 0.00086287 | 0.28881936 | 0.297 | 0.12 | 1 |
| **Ybx1** | 0.00092722 | 0.35350046 | 0.747 | 0.393 | 1 |
| **Pfn1** | 0.00097685 | 0.27430404 | 0.758 | 0.41 | 1 |
| **Usp15** | 0.00104028 | 0.27858843 | 0.275 | 0.109 | 1 |
| **Atf3** | 0.00104404 | 0.43337359 | 0.945 | 0.732 | 1 |
| **Ppib** | 0.00104682 | 0.31185809 | 0.824 | 0.47 | 1 |
| **Cald1** | 0.0011938 | 0.35854715 | 0.747 | 0.432 | 1 |
| **Snx3** | 0.00119632 | 0.25441633 | 0.791 | 0.475 | 1 |
| **Ccnt1** | 0.00135666 | -0.2619506 | 0.275 | 0.104 | 1 |
| **Sox17** | 0.00140715 | 0.31588943 | 0.319 | 0.142 | 1 |
| **Arpc2** | 0.00154857 | 0.25740478 | 0.681 | 0.372 | 1 |
| **Smdt1** | 0.00155829 | 0.27017754 | 0.945 | 0.514 | 1 |
| **Nr4a2** | 0.00193256 | 0.29376713 | 0.297 | 0.126 | 1 |
| **Gng5** | 0.00206792 | 0.31978187 | 0.967 | 0.727 | 1 |
| **Procr** | 0.00215335 | 0.38128242 | 0.264 | 0.109 | 1 |
| **Ctdsp2** | 0.0021841 | -0.2637125 | 0.374 | 0.164 | 1 |
| **Sat1** | 0.00223796 | 0.34069017 | 0.637 | 0.339 | 1 |
| **Ilkap** | 0.00253157 | -0.272624 | 0.319 | 0.131 | 1 |
| **Lsm8** | 0.00278576 | -0.2659257 | 0.429 | 0.197 | 1 |
| **Atox1** | 0.00326935 | 0.29149182 | 0.989 | 0.852 | 1 |
| **Rpl28** | 0.00327113 | 0.28660835 | 0.945 | 0.749 | 1 |
| **Rnpepl1** | 0.00347133 | -0.2751504 | 0.264 | 0.104 | 1 |
| **Hypk** | 0.00396152 | -0.2581699 | 0.429 | 0.197 | 1 |
| **Ewsr1** | 0.0040526 | -0.2798279 | 0.429 | 0.202 | 1 |
| **Gabarap** | 0.00408194 | 0.29201208 | 0.868 | 0.541 | 1 |
| **Cr1l** | 0.00440365 | -0.2638389 | 0.352 | 0.158 | 1 |
| **Pin1** | 0.00476559 | -0.3028477 | 0.319 | 0.137 | 1 |
| **mt-Nd2** | 0.00481793 | -0.6023989 | 0.989 | 0.885 | 1 |
| **Rbm42** | 0.00538686 | -0.2771329 | 0.341 | 0.153 | 1 |
| **Lrpap1** | 0.0057166 | -0.2728411 | 0.308 | 0.137 | 1 |
| **Lyn** | 0.00585466 | -0.2517597 | 0.527 | 0.268 | 1 |
| **Rab6a** | 0.00590568 | -0.2733225 | 0.275 | 0.12 | 1 |
| **Dctn6** | 0.00614776 | -0.3702317 | 0.363 | 0.169 | 1 |
| **Hsd3b7** | 0.00631825 | -0.3158037 | 0.275 | 0.12 | 1 |
| **Ubr4** | 0.00648139 | -0.2966351 | 0.264 | 0.115 | 1 |
| **Clec14a** | 0.00681089 | -0.2803831 | 0.429 | 0.213 | 1 |
| **Wls** | 0.00741528 | -0.3160307 | 0.473 | 0.235 | 1 |
| **Egln2** | 0.00830094 | -0.2518101 | 0.264 | 0.115 | 1 |
| **Plxna2** | 0.0090196 | -0.2667012 | 0.286 | 0.131 | 1 |
| **Hspa5** | 0.00922785 | 0.33189693 | 0.714 | 0.464 | 1 |
| **Phip** | 0.00931784 | -0.2675513 | 0.275 | 0.12 | 1 |
| **Mga** | 0.00981751 | -0.2505006 | 0.264 | 0.12 | 1 |
| **Ppp1r2** | 0.01002555 | 0.26369571 | 0.626 | 0.377 | 1 |
| **Tmem176a** | 0.01007028 | -0.2547464 | 0.374 | 0.18 | 1 |
| **Acadm** | 0.01018821 | -0.3610654 | 0.352 | 0.169 | 1 |
| **Rgs3** | 0.01047053 | -0.4080941 | 0.297 | 0.137 | 1 |
| **Cldn5** | 0.01107038 | 0.28957366 | 0.901 | 0.65 | 1 |
| **Eid1** | 0.01222236 | -0.2585693 | 0.582 | 0.306 | 1 |
| **Ifitm3** | 0.01271469 | -0.4661341 | 0.989 | 0.902 | 1 |
| **Egfl7** | 0.01293005 | -0.4525999 | 0.901 | 0.743 | 1 |
| **Mrpl30** | 0.01357206 | -0.2731676 | 0.396 | 0.197 | 1 |
| **Snhg12** | 0.0136351 | -0.3071653 | 0.253 | 0.115 | 1 |
| **Rpl3** | 0.01365272 | 0.32861386 | 0.835 | 0.568 | 1 |
| **Smc6** | 0.01380389 | -0.3159959 | 0.374 | 0.186 | 1 |
| **Vdac3** | 0.01388041 | -0.3137591 | 0.516 | 0.273 | 1 |
| **Chchd3** | 0.01422463 | -0.2938596 | 0.253 | 0.115 | 1 |
| **Csnk2b** | 0.01498423 | -0.2821628 | 0.451 | 0.235 | 1 |
| **Ccl21a** | 0.01506638 | -1.1773967 | 0.33 | 0.448 | 1 |
| **Naa38** | 0.01548165 | -0.3494749 | 0.385 | 0.197 | 1 |
| **Egln1** | 0.01557965 | -0.2969127 | 0.253 | 0.115 | 1 |
| **Dab2** | 0.01602555 | -0.2777321 | 0.615 | 0.339 | 1 |
| **Arid5b** | 0.01683624 | -0.2950721 | 0.516 | 0.273 | 1 |
| **Dab2ip** | 0.01726364 | -0.2797188 | 0.352 | 0.18 | 1 |
| **Gimap1** | 0.01785227 | -0.2726385 | 0.352 | 0.186 | 1 |
| **Grpel1** | 0.01827194 | -0.2608677 | 0.451 | 0.24 | 1 |
| **Gatad2b** | 0.01896613 | -0.4116625 | 0.286 | 0.137 | 1 |
| **Eif3d** | 0.01896613 | -0.3245184 | 0.286 | 0.137 | 1 |
| **Usp10** | 0.01897547 | -0.2836486 | 0.253 | 0.12 | 1 |
| **Sin3b** | 0.01942285 | -0.2744892 | 0.363 | 0.186 | 1 |
| **Ptpn18** | 0.01975677 | -0.2610199 | 0.396 | 0.213 | 1 |
| **Cenpx** | 0.01977609 | -0.3508775 | 0.462 | 0.251 | 1 |
| **Smim10l1** | 0.0198616 | -0.2756024 | 0.352 | 0.18 | 1 |
| **Ptn** | 0.0199237 | 0.43723825 | 0.286 | 0.158 | 1 |
| **Hmgb2** | 0.0200078 | -0.317528 | 0.275 | 0.131 | 1 |
| **Copz2** | 0.02161766 | -0.2520767 | 0.264 | 0.126 | 1 |
| **Pink1** | 0.02256172 | -0.3398852 | 0.253 | 0.12 | 1 |
| **Cst3** | 0.02268239 | -0.4492529 | 0.945 | 0.792 | 1 |
| **Arglu1** | 0.02323602 | -0.2804207 | 0.44 | 0.23 | 1 |
| **Fkbp2** | 0.02337173 | -0.3800924 | 0.44 | 0.235 | 1 |
| **Sys1** | 0.02410894 | -0.271555 | 0.451 | 0.24 | 1 |
| **Pttg1** | 0.0241357 | -0.265476 | 0.374 | 0.191 | 1 |
| **Rgl2** | 0.02480145 | -0.3459866 | 0.297 | 0.153 | 1 |
| **Kidins220** | 0.02523303 | -0.291756 | 0.253 | 0.126 | 1 |
| **Usp9x** | 0.02625791 | -0.2759278 | 0.264 | 0.131 | 1 |
| **Ptov1** | 0.0269251 | -0.4373089 | 0.264 | 0.131 | 1 |
| **Pofut2** | 0.02709417 | -0.3504275 | 0.264 | 0.131 | 1 |
| **Elmo1** | 0.02753645 | -0.286755 | 0.33 | 0.175 | 1 |
| **Psmb8** | 0.02826527 | -0.3654584 | 0.374 | 0.197 | 1 |
| **Mif** | 0.02846786 | -0.2996378 | 0.319 | 0.164 | 1 |
| **Hsbp1** | 0.02907498 | -0.260428 | 0.615 | 0.339 | 1 |
| **Tgfbr3** | 0.02944374 | -0.2872485 | 0.275 | 0.137 | 1 |
| **Ier5l** | 0.03047902 | -0.421973 | 0.418 | 0.224 | 1 |
| **Pgp** | 0.03071573 | -0.3239537 | 0.253 | 0.126 | 1 |
| **Notch1** | 0.03080053 | -0.2624507 | 0.495 | 0.273 | 1 |
| **Txndc12** | 0.03121835 | -0.2893367 | 0.308 | 0.164 | 1 |
| **Sh2d3c** | 0.03266746 | -0.3145852 | 0.308 | 0.164 | 1 |
| **Gm26532** | 0.0341145 | -0.360657 | 0.714 | 0.432 | 1 |
| **Brd9** | 0.03532281 | -0.3533592 | 0.429 | 0.235 | 1 |
| **Neat1** | 0.03532665 | -0.3263718 | 0.549 | 0.306 | 1 |
| **Tnks2** | 0.03565407 | -0.2515555 | 0.264 | 0.137 | 1 |
| **Ndufv3** | 0.03748379 | -0.3406088 | 0.626 | 0.35 | 1 |
| **Ctsz** | 0.0382034 | -0.2613185 | 0.396 | 0.23 | 1 |
| **Cfh** | 0.04039623 | -0.3330194 | 0.418 | 0.24 | 1 |
| **Tnrc6c** | 0.04273448 | -0.2750424 | 0.44 | 0.246 | 1 |
| **Tiparp** | 0.04312745 | -0.2504063 | 0.264 | 0.142 | 1 |
| **Tomm22** | 0.04344234 | -0.2908938 | 0.516 | 0.29 | 1 |
| **Rpp21** | 0.04360576 | -0.4297338 | 0.319 | 0.169 | 1 |
| **Tmbim4** | 0.04363604 | -0.4122243 | 0.396 | 0.219 | 1 |
| **Tma7** | 0.04449075 | -0.2529658 | 0.538 | 0.306 | 1 |
| **Fnta** | 0.04536867 | -0.282026 | 0.297 | 0.158 | 1 |
| **Lamc1** | 0.04688413 | -0.2864634 | 0.275 | 0.148 | 1 |
| **Degs1** | 0.04790815 | -0.4443358 | 0.418 | 0.23 | 1 |
| **Saraf** | 0.04903187 | -0.2789539 | 0.352 | 0.202 | 1 |
| **Mrps18c** | 0.04965982 | -0.335877 | 0.396 | 0.23 | 1 |
| **Adrm1** | 0.05412524 | -0.339614 | 0.275 | 0.148 | 1 |
| **Kdm6b** | 0.05527355 | -0.2570504 | 0.297 | 0.164 | 1 |
| **Kmt2a** | 0.05529435 | -0.3852326 | 0.341 | 0.186 | 1 |
| **Cggbp1** | 0.0561949 | -0.2817413 | 0.527 | 0.311 | 1 |
| **Fam43a** | 0.05782143 | -0.2843158 | 0.418 | 0.246 | 1 |
| **Tcf25** | 0.05996489 | -0.2578353 | 0.516 | 0.306 | 1 |
| **Adgrf5** | 0.06002101 | -0.3484318 | 0.516 | 0.295 | 1 |
| **Slco3a1** | 0.06045013 | -0.3492579 | 0.396 | 0.23 | 1 |
| **Atp5g3** | 0.0622568 | -0.3950906 | 0.637 | 0.355 | 1 |
| **Ypel3** | 0.06244891 | -0.3555447 | 0.473 | 0.273 | 1 |
| **Etnk1** | 0.06303954 | -0.3297757 | 0.33 | 0.186 | 1 |
| **mt-Nd4l** | 0.06388328 | -0.6221701 | 0.923 | 0.787 | 1 |
| **Tcea1** | 0.06579909 | -0.3168662 | 0.538 | 0.311 | 1 |
| **Ggnbp2** | 0.06703358 | -0.3265603 | 0.275 | 0.153 | 1 |
| **H1f0** | 0.06773695 | -0.3422277 | 0.275 | 0.153 | 1 |
| **Psenen** | 0.06887748 | -0.2695476 | 0.56 | 0.333 | 1 |
| **Ddit4** | 0.07001983 | -0.3332343 | 0.286 | 0.158 | 1 |
| **Tle4** | 0.07179114 | -0.4802766 | 0.275 | 0.148 | 1 |
| **Ndufs6** | 0.07498102 | -0.3264791 | 0.516 | 0.306 | 1 |
| **Pcsk6** | 0.07658631 | -0.3712171 | 0.352 | 0.202 | 1 |
| **Vps28** | 0.07776062 | -0.3543961 | 0.505 | 0.29 | 1 |
| **Vkorc1** | 0.07783013 | -0.3011844 | 0.286 | 0.164 | 1 |
| **Tra2a** | 0.07941154 | -0.296453 | 0.407 | 0.24 | 1 |
| **Ndufc1** | 0.08448555 | -0.2983084 | 0.736 | 0.432 | 1 |
| **Ccdc85b** | 0.0849134 | -0.2569683 | 0.626 | 0.372 | 1 |
| **Nid1** | 0.08584796 | -0.3444762 | 0.275 | 0.158 | 1 |
| **Ndufa11** | 0.08611427 | -0.316741 | 0.659 | 0.388 | 1 |
| **Psmd1** | 0.08642727 | -0.3160835 | 0.374 | 0.219 | 1 |
| **Sdhaf4** | 0.08651987 | -0.3493589 | 0.275 | 0.153 | 1 |
| **Sipa1** | 0.08930333 | -0.4894227 | 0.286 | 0.164 | 1 |
| **Nktr** | 0.09154979 | -0.3799742 | 0.308 | 0.18 | 1 |
| **Cav2** | 0.09238713 | -0.303259 | 0.538 | 0.311 | 1 |
| **Eci1** | 0.09516216 | -0.4485073 | 0.275 | 0.158 | 1 |
| **Il6st** | 0.09558264 | -0.5767679 | 0.374 | 0.213 | 1 |
| **Kbtbd11** | 0.09605785 | -0.4344783 | 0.264 | 0.148 | 1 |
| **Ctsb** | 0.0975171 | -0.272268 | 0.527 | 0.333 | 1 |
| **Baz2b** | 0.10180937 | -0.3797874 | 0.275 | 0.158 | 1 |
| **Lars2** | 0.10461848 | -1.625086 | 0.549 | 0.514 | 1 |
| **Nudcd3** | 0.10715969 | -0.3366264 | 0.429 | 0.262 | 1 |
| **Kif2a** | 0.10780029 | -0.2774292 | 0.275 | 0.164 | 1 |
| **Mtdh** | 0.11126876 | -0.4326722 | 0.516 | 0.322 | 1 |
| **Pkhd1l1** | 0.11212466 | -0.255185 | 0.451 | 0.284 | 1 |
| **2-Mar** | 0.11237918 | -0.3184093 | 0.363 | 0.219 | 1 |
| **Psme1** | 0.11244531 | -0.4359715 | 0.637 | 0.372 | 1 |
| **Mrps24** | 0.11255756 | -0.2900752 | 0.593 | 0.361 | 1 |
| **Bsg** | 0.11335012 | -0.7653775 | 0.538 | 0.475 | 1 |
| **Cisd1** | 0.11840743 | -0.380575 | 0.297 | 0.18 | 1 |
| **R3hdm2** | 0.11880331 | -0.3394536 | 0.253 | 0.148 | 1 |
| **Tnfsf12** | 0.12020539 | -0.2963932 | 0.286 | 0.169 | 1 |
| **2310001H17Rik** | 0.12267451 | -0.4433356 | 0.286 | 0.169 | 1 |
| **Snrnp70** | 0.12435472 | -0.3915136 | 0.407 | 0.251 | 1 |
| **Plk2** | 0.12572659 | -0.333685 | 0.659 | 0.41 | 1 |
| **Ccdc59** | 0.12618281 | -0.479924 | 0.363 | 0.219 | 1 |
| **Emd** | 0.12695824 | -0.3196561 | 0.264 | 0.158 | 1 |
| **Snhg8** | 0.12998745 | -0.3195773 | 0.253 | 0.148 | 1 |
| **Pgls** | 0.1323148 | -0.3780526 | 0.385 | 0.23 | 1 |
| **Dek** | 0.13909947 | -0.271774 | 0.396 | 0.246 | 1 |
| **Krtcap2** | 0.13913564 | -0.3942641 | 0.681 | 0.415 | 1 |
| **Ets2** | 0.14347838 | -0.3780994 | 0.451 | 0.29 | 1 |
| **Fkbp8** | 0.14501069 | -0.3543831 | 0.385 | 0.24 | 1 |
| **Hnrnpl** | 0.1471978 | -0.4237636 | 0.363 | 0.224 | 1 |
| **Tmem234** | 0.14772011 | -0.4597583 | 0.582 | 0.344 | 1 |
| **Fgfr1op2** | 0.14915318 | -0.3854715 | 0.253 | 0.153 | 1 |
| **Dock9** | 0.14933048 | -0.3323732 | 0.56 | 0.35 | 1 |
| **Bptf** | 0.15150768 | -0.3419462 | 0.341 | 0.213 | 1 |
| **Tanc1** | 0.1520406 | -0.3346106 | 0.363 | 0.23 | 1 |
| **Rtl8b** | 0.15368655 | -0.2575893 | 0.253 | 0.158 | 1 |
| **Commd6** | 0.16204118 | -0.2555473 | 0.253 | 0.158 | 1 |
| **Galnt18** | 0.16396843 | -0.3783377 | 0.407 | 0.257 | 1 |
| **Herc1** | 0.16603467 | -0.3856614 | 0.275 | 0.175 | 1 |
| **Atf4** | 0.16735363 | -0.472917 | 0.495 | 0.295 | 1 |
| **Tsg101** | 0.17237282 | -0.4558733 | 0.264 | 0.164 | 1 |
| **Ndufb4** | 0.17804189 | -0.2663111 | 0.725 | 0.443 | 1 |
| **Scand1** | 0.17937972 | -0.4211356 | 0.44 | 0.279 | 1 |
| **Arhgap23** | 0.18054064 | -0.3023986 | 0.253 | 0.158 | 1 |
| **Selenow** | 0.18147467 | -0.3632395 | 0.549 | 0.344 | 1 |
| **Hmgb1** | 0.18719151 | -0.2824717 | 0.879 | 0.732 | 1 |
| **Ushbp1** | 0.19331838 | -0.5673486 | 0.374 | 0.235 | 1 |
| **Cnot3** | 0.21247305 | -0.2674994 | 0.253 | 0.164 | 1 |
| **Dnajb1** | 0.21815416 | -0.5572085 | 0.385 | 0.246 | 1 |
| **Gsto1** | 0.22026923 | -0.4682689 | 0.308 | 0.197 | 1 |
| **Dock6** | 0.22452175 | -0.4729865 | 0.407 | 0.262 | 1 |
| **Srsf2** | 0.22872594 | -0.3263281 | 0.516 | 0.328 | 1 |
| **U2af1** | 0.22902709 | -0.3229454 | 0.484 | 0.322 | 1 |
| **Mrpl48** | 0.23066013 | -0.4712419 | 0.253 | 0.164 | 1 |
| **Cebpb** | 0.23326929 | -0.64765 | 0.44 | 0.279 | 1 |
| **Igfbp7** | 0.23511879 | -1.0542259 | 0.527 | 0.459 | 1 |
| **Btg2** | 0.2387057 | -0.340589 | 0.736 | 0.464 | 1 |
| **Grcc10** | 0.24686083 | -0.2916057 | 0.549 | 0.372 | 1 |
| **Tmem176b** | 0.25237509 | -0.2660778 | 0.385 | 0.262 | 1 |
| **Ndufb7** | 0.25598559 | -0.4536479 | 0.549 | 0.35 | 1 |
| **Tmsb10** | 0.257824 | -0.2566069 | 1 | 0.934 | 1 |
| **Ppp2r5c** | 0.26577884 | -0.3816226 | 0.275 | 0.18 | 1 |
| **Flt4** | 0.27381111 | -0.3282753 | 0.549 | 0.35 | 1 |
| **Robo4** | 0.27493679 | -0.4156572 | 0.308 | 0.202 | 1 |
| **Srrm2** | 0.27680313 | -0.3295627 | 0.736 | 0.475 | 1 |
| **B230219D22Rik** | 0.29390329 | -0.3309439 | 0.637 | 0.393 | 1 |
| **Mmrn1** | 0.2946764 | -0.5810928 | 0.484 | 0.311 | 1 |
| **Trmt112** | 0.2974565 | -0.3993115 | 0.396 | 0.262 | 1 |
| **Cbfa2t3** | 0.30966557 | -0.4617713 | 0.286 | 0.191 | 1 |
| **Tns1** | 0.32081459 | -0.4456551 | 0.495 | 0.333 | 1 |
| **Hist1h2bc** | 0.32190865 | -0.3711579 | 0.253 | 0.169 | 1 |
| **Ppp1cc** | 0.32301954 | -0.3731566 | 0.462 | 0.311 | 1 |
| **Ndufa5** | 0.32328003 | -0.2954596 | 0.736 | 0.475 | 1 |
| **Palm** | 0.32498488 | -0.393451 | 0.286 | 0.197 | 1 |
| **Fermt2** | 0.33164136 | -0.3544613 | 0.648 | 0.426 | 1 |
| **Pop5** | 0.3450829 | -0.3882946 | 0.297 | 0.208 | 1 |
| **Lama4** | 0.35765492 | -0.2696551 | 0.582 | 0.41 | 1 |
| **Chic2** | 0.36139491 | -0.3431842 | 0.33 | 0.23 | 1 |
| **Srsf11** | 0.37235715 | -0.4094175 | 0.44 | 0.301 | 1 |
| **Sned1** | 0.37814826 | -0.4095908 | 0.473 | 0.333 | 1 |
| **Cd63** | 0.38828516 | -0.2678592 | 0.681 | 0.443 | 1 |
| **Tgfbr2** | 0.39021778 | -0.447202 | 0.626 | 0.388 | 1 |
| **Ndufb10** | 0.42651619 | -0.4354362 | 0.604 | 0.383 | 1 |
| **Atp5k** | 0.44989608 | -0.4389781 | 0.692 | 0.464 | 1 |
| **Gm47283** | 0.46803317 | -0.7131728 | 0.538 | 0.432 | 1 |
| **Cebpd** | 0.48084284 | -0.2936667 | 0.912 | 0.667 | 1 |
| **Cox6c** | 0.50186653 | -0.2546826 | 0.978 | 0.831 | 1 |
| **Pard6g** | 0.50232653 | -0.5799683 | 0.516 | 0.361 | 1 |
| **Pim3** | 0.50344598 | -0.5170018 | 0.462 | 0.317 | 1 |
| **Gadd45g** | 0.50471765 | -0.5964558 | 0.484 | 0.339 | 1 |
| **mt-Atp8** | 0.52842917 | -0.6502659 | 0.791 | 0.601 | 1 |
| **mt-Nd5** | 0.5367032 | -0.5060443 | 0.846 | 0.601 | 1 |
| **Bst2** | 0.5367136 | -0.6504045 | 0.637 | 0.432 | 1 |
| **Nop53** | 0.54982756 | -0.5096591 | 0.319 | 0.235 | 1 |
| **Myl12b** | 0.56342855 | -0.5378612 | 0.758 | 0.525 | 1 |
| **Plvap** | 0.57618679 | -0.2763148 | 0.637 | 0.443 | 1 |
| **Cyba** | 0.57859156 | -0.5515915 | 0.484 | 0.339 | 1 |
| **Lrg1** | 0.61196271 | -0.5436858 | 0.681 | 0.481 | 1 |
| **Cox4i1** | 0.61420842 | -0.3168232 | 0.879 | 0.623 | 1 |
| **Tm4sf1** | 0.62518037 | -0.3898934 | 0.912 | 0.738 | 1 |
| **Fosb** | 0.62797891 | -0.3816809 | 0.813 | 0.601 | 1 |
| **Clec2d** | 0.63335451 | -0.4822726 | 0.615 | 0.481 | 1 |
| **AY036118** | 0.6359889 | -0.7034626 | 0.385 | 0.306 | 1 |
| **Egr1** | 0.66647252 | -0.5131691 | 0.681 | 0.546 | 1 |
| **Ltbp4** | 0.67265293 | -0.2898987 | 0.846 | 0.59 | 1 |
| **Chmp2a** | 0.67552568 | -0.3202394 | 0.692 | 0.47 | 1 |
| **Ugcg** | 0.70811872 | -0.5047841 | 0.33 | 0.251 | 1 |
| **Pabpc1** | 0.7689826 | -0.3449492 | 0.615 | 0.459 | 1 |
| **Mtch1** | 0.80513824 | -0.4905542 | 0.637 | 0.443 | 1 |
| **Kank3** | 0.80809582 | -0.4703469 | 0.67 | 0.459 | 1 |
| **Apoe** | 0.84499679 | -0.6661397 | 0.308 | 0.24 | 1 |
| **Gpm6a** | 0.85426117 | -0.5030455 | 0.429 | 0.355 | 1 |
| **Ndufs5** | 0.86921439 | -0.3922484 | 0.637 | 0.464 | 1 |
| **Mt1** | 0.8750415 | -0.4712849 | 0.33 | 0.29 | 1 |
| **Slc25a4** | 0.87555902 | -0.3165486 | 0.714 | 0.508 | 1 |
| **Ctsd** | 0.87918902 | -0.6974264 | 0.418 | 0.328 | 1 |
| **Rpl13a** | 0.88664816 | -0.3958025 | 0.714 | 0.53 | 1 |
| **Psmd8** | 0.90986757 | -0.7896764 | 0.396 | 0.311 | 1 |
| **Ndufa3** | 0.91259101 | -0.6750198 | 0.67 | 0.448 | 1 |
| **Map1lc3b** | 0.9306778 | -0.3424144 | 0.637 | 0.448 | 1 |
| **Ppp2r5a** | 0.95211462 | -0.557848 | 0.527 | 0.383 | 1 |
| **Slc3a2** | 0.96226408 | -0.5787949 | 0.44 | 0.35 | 1 |
