## Supplemental figures for "Transcriptomic profiling of Schlemm’s canal cells reveals a lymphatic-biased identity and three major cell states"

Supporting Figures

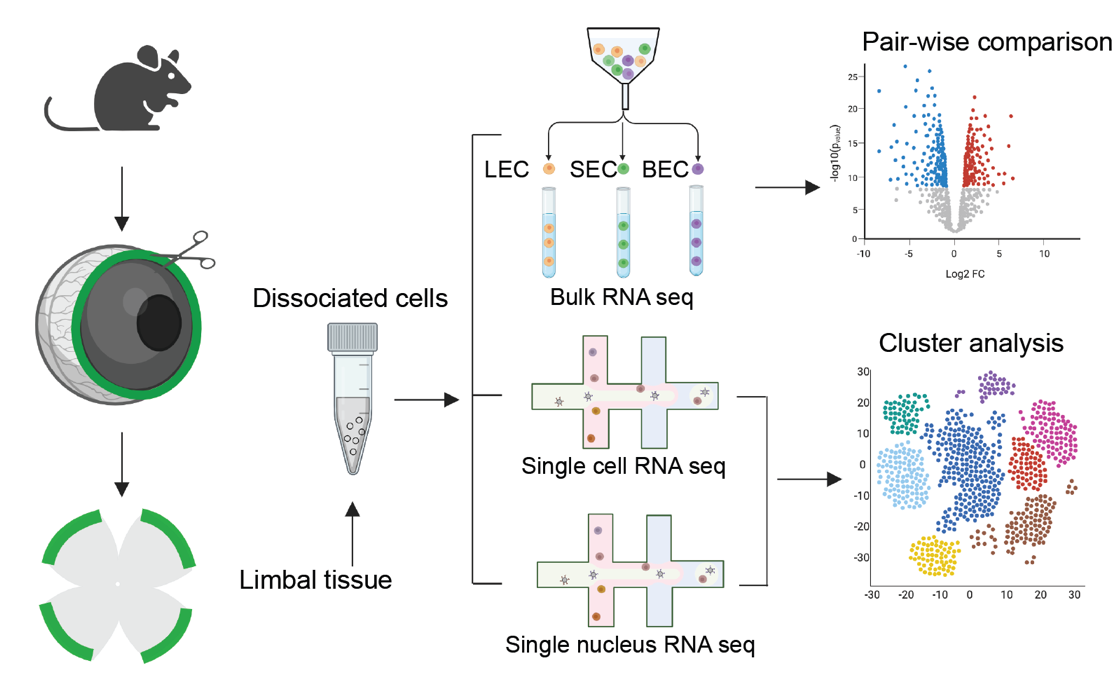

**Fig. S1.** **RNA sequencing of anterior segment limbal tissue.** Schematic showing tissue processing for bulk, single cell, and single nucleus RNA sequencing methods. LEC: Lymphatic endothelial cells, SEC: Schlemm’s canal endothelial cells, BEC: Blood endothelial cells. Created with BioRender.com.

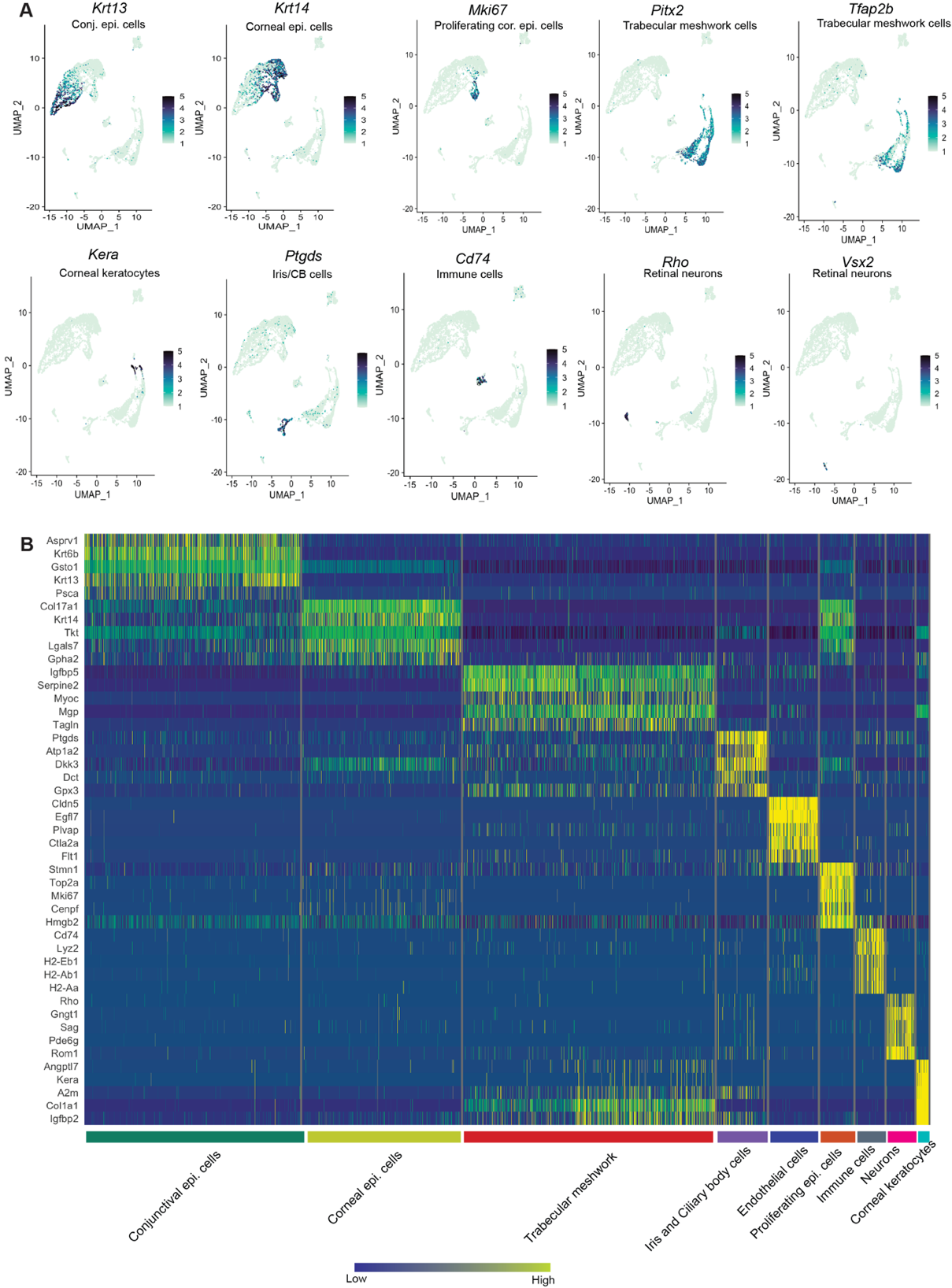

**Fig. S2. Cell types in mouse limbal tissue.** A. Expression levels of various genes previously identified to be specific to individual cell types on a UMAP rendering of single cell clusters. Epithelial cells comprised of conjunctival (*Krt13*), corneal (*Krt14*), and proliferating corneal epithelial cells (*Mki67*); neural crest or periocular mesenchyme derived cells (*Pitx2*) comprised of trabecular meshwork cells (*Tfap2b*); corneal keratocytes (*Kera*); ciliary body (CB) and iris cells (*Ptgds*); immune cells (*Cd74*); and neurons (*Vsx2*, *Rho*) were identified (Figure S2). B. Heatmap of differentially expressed genes across various cell clusters identified in the limbal tissue.

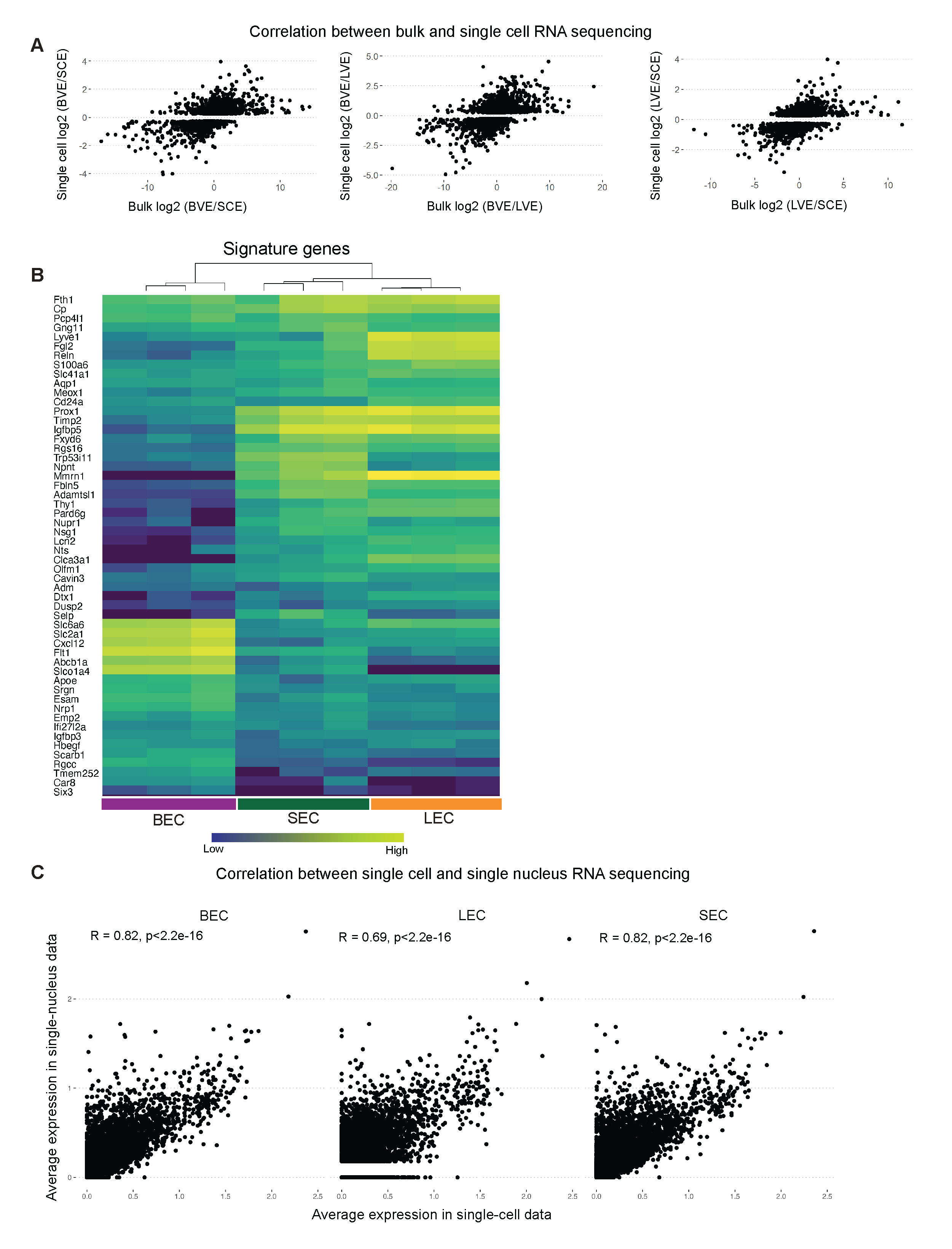

**Fig. S3: Correlation between sequencing modalities**. A. Correlation between bulk and scRNA-seq comparing log of gene expression ratio between endothelial cell types. B. Hierarchical clustering of signature genes obtained from scRNA-seq in 2F using bulk RNA sequencing data. C. Correlation between sc and snRNA-seq comparing average gene expression across endothelial cell types. BEC: Blood endothelial cell, LEC: Lymphatic endothelial cell, SEC: Schlemm’s canal endothelial cell.

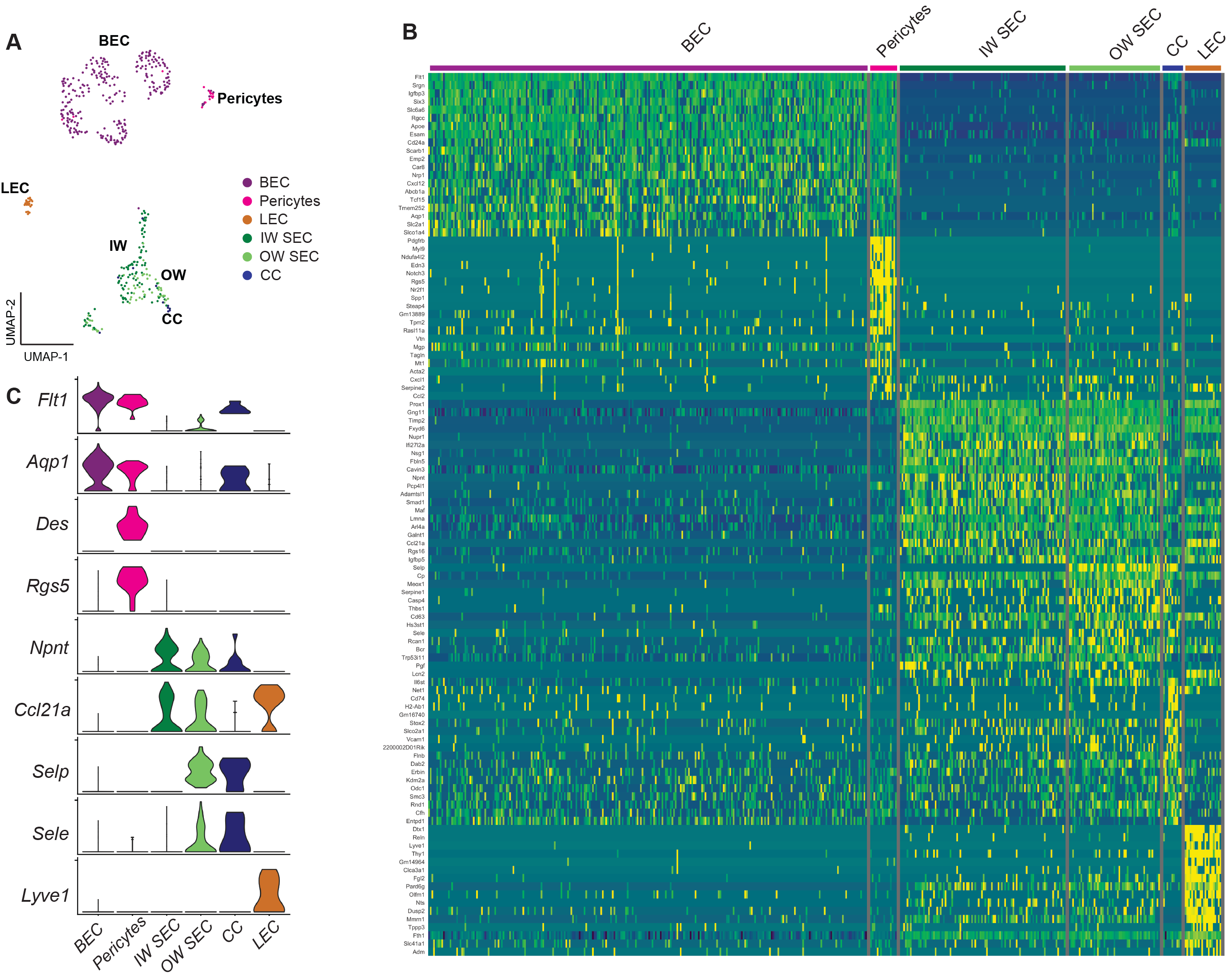

**Fig. S4. EC and pericyte transcriptomes in scRNA-seq data guided by integrated multimodal data.** A. Manually re-clustered UMAP showing BECs, pericytes, LECs, IW SECs, OW SECs, and CCs. B, C. Differences in levels of gene expression among BEC, pericytes, LEC, IW SEC, OW SEC, and CC expressed in a heatmap (B) and a violin plot (C). OW: Outer wall, IW: Inner wall, CC: Collector channels, BEC: Blood endothelial cell, LEC: Lymphatic endothelial cell, SEC: Schlemm’s canal endothelial cell.

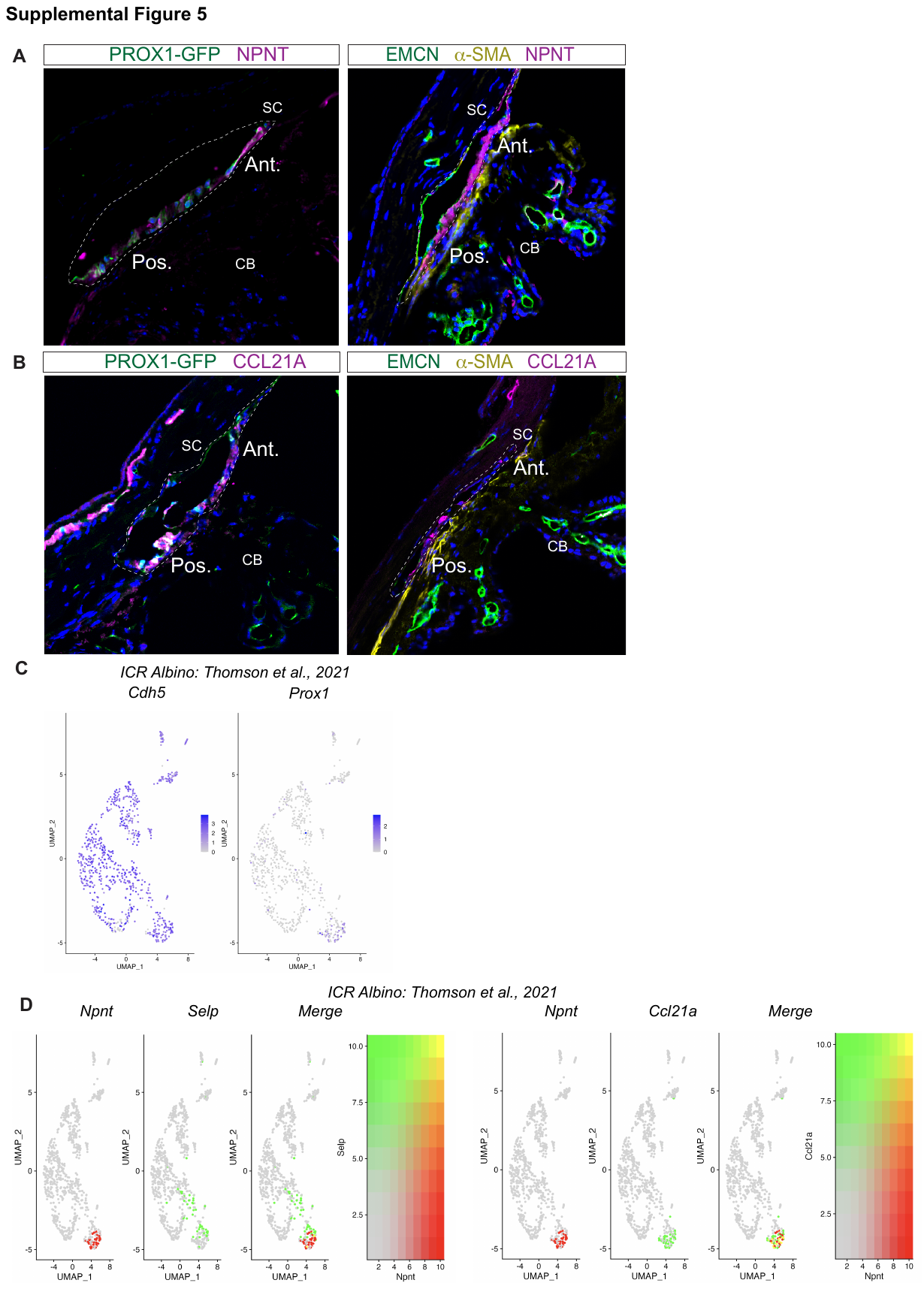

**Fig. S5: Biased but variable localization of NPNT and CCL21A in IW of SC.** A. Immunofluorescence (IF) of NPNT shows variability in expression with a gradient of high expression in the anterior portion of SC (left panel) and uniform expression throughout IW of SC (middle panel) in flash-frozen sections. In-situ hybridization using RNAscope® shows the expression of *Npnt* in IW with an anterior expression bias (left panel) B. IF of CCL21A shows variability in expression with a gradient of high expression in the posterior portion of SC (right panel) and uniform expression throughout IW of SC (left panel) in flash-frozen sections. DAPI in blue labels nuclei in all panels. C, D. Gene expression analysis of endothelial cell subset in published dataset (Thomson et al., 2021) shows similar segregation of *Npnt, Selp,* and *Ccl21a* expression in SECs as seen in our dataset. CB: ciliary body, SC: Schlemm’s canal. Ant.: Anterior Schlemm’s canal, Post.: Posterior Schlemm’s canal.

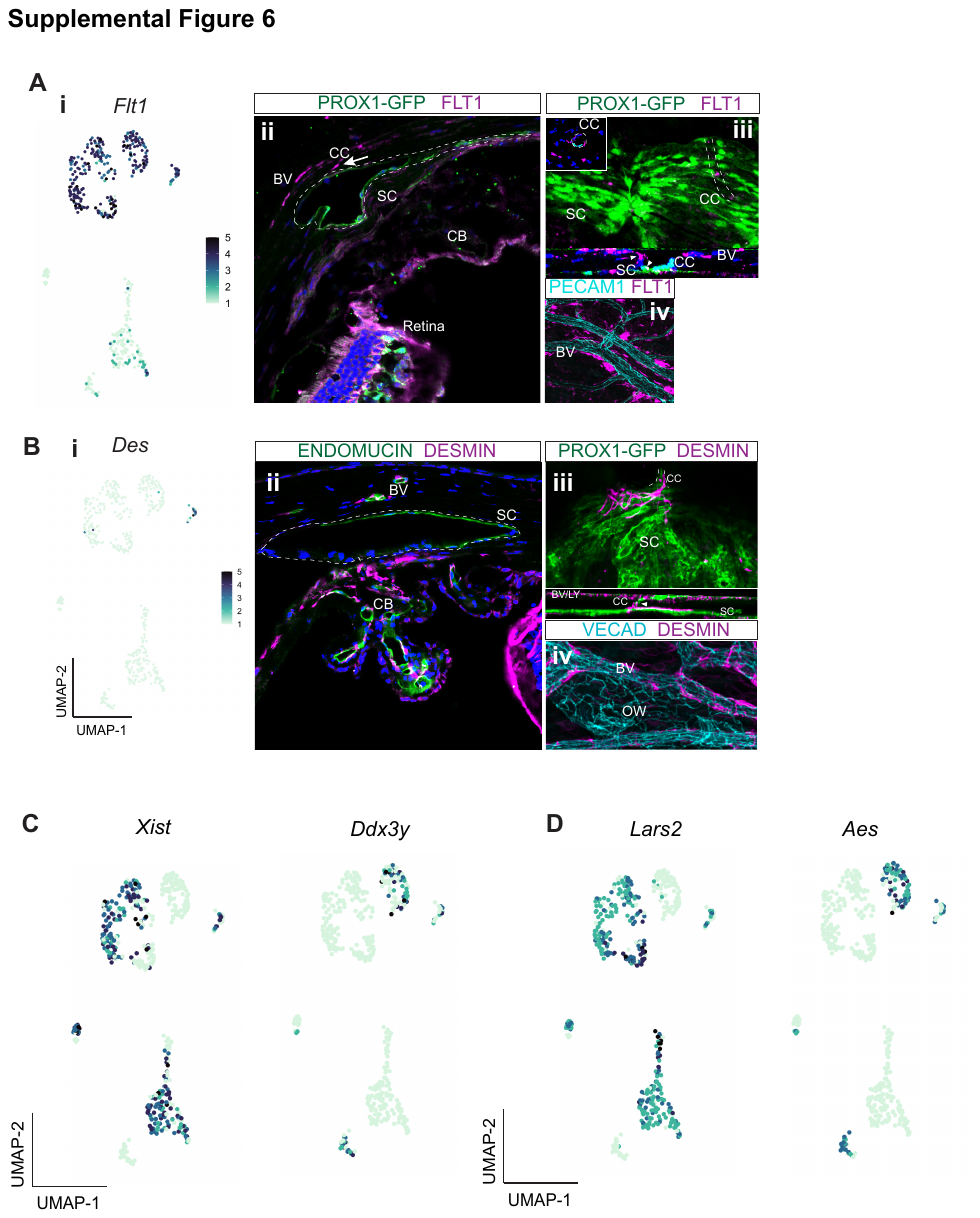

**Fig. S6:** **Collector channels, pericytes, and sex-dependent differences within the major EC cluster.** A. Flt1 is expressed primarily in BECs and in a small subset of SECs in scRNA-seq analysis (i) and corresponding immunofluorescence (IF) of FLT1 in frozen section (ii) and whole-mount (iii-iv). B. *Des* is expressed in pericytes, a subgroup of BECs in scRNA-seq analysis (i). Corresponding IF analysis shows its expression in pericytes along collector channels (ii, frozen section, iii-iv whole mount). DAPI in blue labels nuclei in all panels. C. Sub-clustering of SECs and sex-specificity of endothelial cells. Sex-specific gene expression of Xist, Ddx3y, Lars2, Aes in ‘male’ and ‘female’ clusters of endothelial cells. CC: Collector channel, CB: Ciliary body, OW: Outer wall, BV: Blood vessel, SC: Schlemm’s canal.

**Fig. S7: Main BEC types within major EC cluster and immunofluorescence confirmation of greater lymphatic polarization of IW.** A. UMAP rendering of BECs subclustered into arteries, capillaries, and veins. B, C. Violin plot and heatmap showing differential gene expression between arteries, capillaries, and veins in limbal BECs. D. *Flt4* is expressed at mid or low levels in SECs and BECs in scRNA-seq analysis (i) and corresponding IF of FLT4 (ii frozen section, iii-iv whole mount). DAPI in blue labels nuclei. BV: Blood vessel, SC: Schlemm’s canal, CB: Ciliary body, TM: Trabecular meshwork, IW: Inner wall.

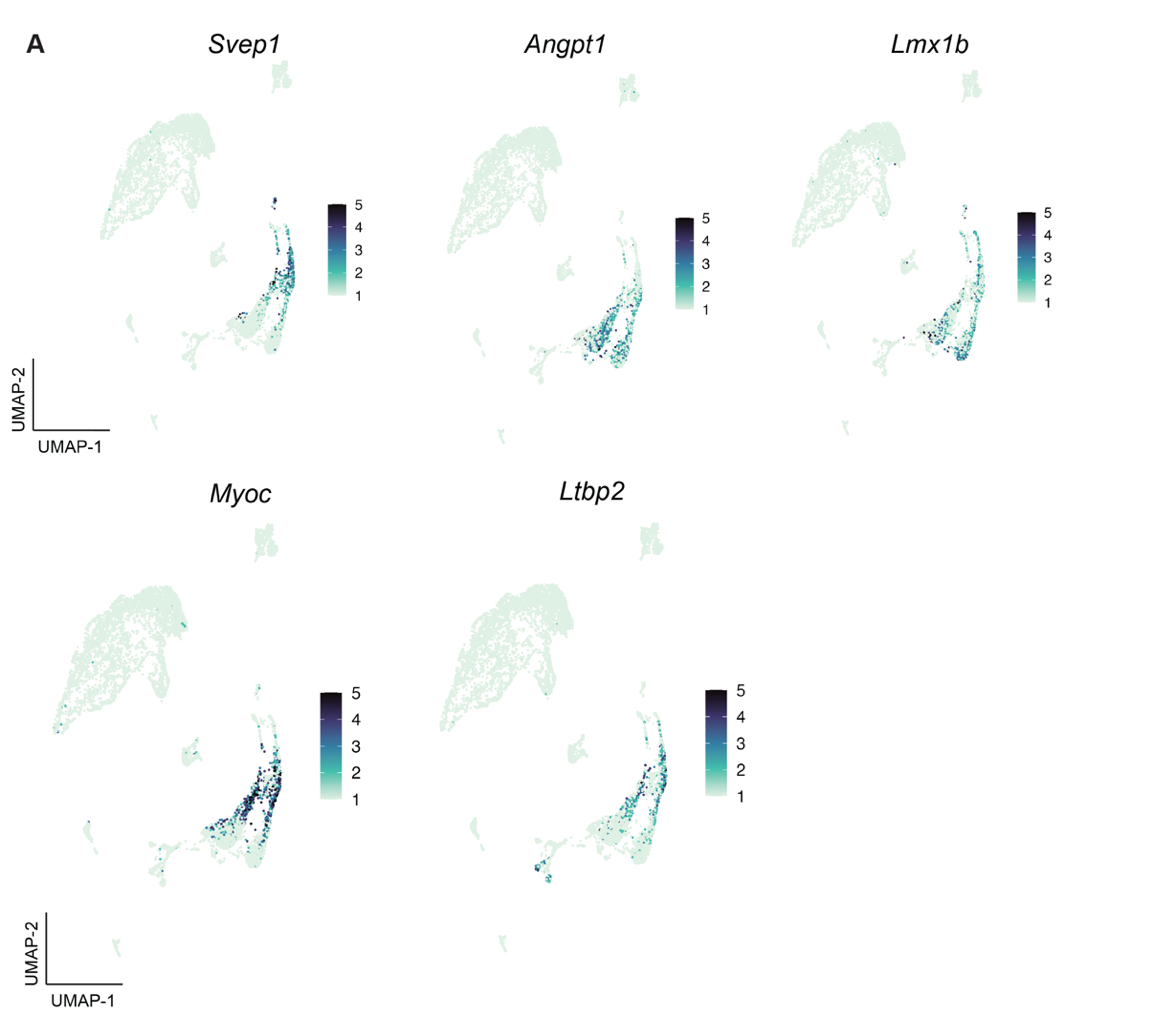

**Fig. S8: Cell type assignment of high IOP and glaucoma genes.** Expression patterns of *Svep1*, *Angpt1*, *Lmx1b*, *Myoc*, and *Ltbp2* in the TM cluster.

**Fig. S9: Gene ontology enrichment analysis highlights key cellular component/ pathways in SECs.** A-C. C-Net plot showing network between individual genes in enriched pathways identified by gene ontology analysis.

**Fig. S10:** **Gene ontology enrichment analysis highlights key pathways in IW and OW SECs**. A-B. Molecular function pathway genes enriched in IW and OW SECs and corresponding c-net plot showing network between individual genes in the pathways.

**Supporting information**

| Antibody | Dilution | Source | ID |
| --- | --- | --- | --- |
| α-SMA | WM (1:50) Sections (1:200) | Abcam | ab5694 |
| CCL21A | WM (1:25)  Sections (1:200) | R&D Systems | AF457-SP |
| DARC/ACKR1 | WM (1:50)  Sections (1:200) | Invitrogen | PA5-47861 |
| Desmin | WM (1:50)  Sections (1:200) | Invitrogen (Thermo Fisher Scientific) | MA532068 |
| Endomucin | WM (1:25)  Sections (1:200) | EBioscience/Invitrogen (Thermo Fisher Scientific) | 14-5851-82 |
| FLT1 | WM (1:50)  Sections (1:100) | Abcam | ab32152 |
| FLT4 | WM (1:50)  Sections (1:200) | R&D Systems | AF743 |
| LYVE1 | WM (1:50)  Sections (1:200) | EBioscience/Invitrogen (Thermo Fisher Scientific) | 14-0443-80 |
| NPNT | WM (1:50)  Sections (1:200) | R&D Systems | AF4298 |
| PECAM (CD31) | WM (1:50)  Sections (1:200) | BD Pharmingen | 550274 |
| SELP | WM (1:50)  Sections (1:100) | R&D Systems | AF737-SP |
| VECAD | WM (1:50)  Sections (1:200) | R&D Systems | AF1002 |

**Table S1. List of antibodies used in the study**

**Table S2. List of other resources and reagents**

| Resource | Source | ID |
| --- | --- | --- |
| Animals | | |
| 129/Sj mice* | Internal | Ref PMID: 33462143 |
| C57BL/6J mice | Jackson Labs | IMSR_JAX:000664 |
| Prox1-GFP BAC transgenic mice (Tg(Prox1-EGFP)KY221Gsat/Mmcd) | MMRRC | MMRRC_031006-UCD |
| Reagents/Chemicals/Other | | |
| 4% PFA | Thermo Fisher Scientific | Cat no. 50-980-495 |
| 40 and 100 μm Falcon cell strainers | Thermo Fisher Scientific | Cat no. 08-771-1 / 08-771-19 |
| Bovine Serum Albumin | Thermo Fisher Scientific | Cat no. AM2616 |
| Collagenase Type 4 | Worthington Biochemical | Cat no. LS004188 |
| DAPI | Thermo Fisher Scientific | Cat no. 62248 |
| Dulbecco's Modified Eagle Medium (DMEM) | Thermo Fisher Scientific | Cat no. 11965-118 |
| Dulbecco's Phosphate Buffered Saline (PBS) | Sigma Aldrich | Cat no. D8662 |
| Earle’s balanced salt solution (EBSS) | Thermo Fisher Scientific | Cat no. 24010-043 |
| Papain Dissociation System | Worthington Biochemical | Cat no. LK003153 |
| Propidium iodide | Thermo Fisher Scientific | Cat no. P1304MP |
| SYTOX green Nucleic Acid Stain | Thermo Fisher Scientific | Cat no. S7020 |

* Genotyping evidence suggests that our 129/Sj strain contains large genomic regions aligning to both 129S6/SvEvTac and 129S1/SvImJ, but is genetically distinct from any other 129 sub-strains.
