## Supplementary material for "Transcriptomic profiling of Schlemm’s canal cells reveals a lymphatic-biased identity and three major cell states": Table 4

| **Sequencing modality** | **Age** | **Total number sequenced** | **Number of ECs sequenced** | **Number of SECs sequenced** |
| --- | --- | --- | --- | --- |
| **Balasubramanian, Kizhatil, et al. 2024** | |  |  |  |
| Single cell C57BL/6J | 3 months | 9272 | 469 | 166 |
| Single nuc C57BL/6J | 3 months | 10,764 | 1287 | 375 |
| Single cell 129/Sj | 2 months | 8812 | 1023 | 362 |
| Total |  | 28848 | 2779 | 903 |
| **Van Zyl et al. 2020** |  |  |  |  |
| Single cell CD1 (albino) | 12 weeks | 5067 | 93 | 25 |
| **Thomson et al. 2021** |  |  |  |  |
| Single cell ICR (albino) | 6 weeks | 19,236 | 763 | 90 |
